# Development of PROTACs for targeted degradation of oncogenic TRK fusions

**DOI:** 10.1101/2025.06.18.660465

**Authors:** Saurav Kumar, Jiewei Jiang, Mia S. Donald-Paladino, Joy Chen, Andrea Gutierrez, Alexander J. Federation, Frank Szulzewsky, Eric C. Holland, Fleur M. Ferguson, Behnam Nabet

**Author notes:** **Correspondence should be addressed to:** Behnam Nabet, Human Biology Division, Fred Hutchinson Cancer Center, 1100 Fairview Avenue N., Seattle, WA, 98109, USA., Fleur M. Ferguson, Department of Chemistry and Biochemistry, University of California San Diego, La Jolla, CA, 92093, USA. Equal contribution (Saurav Kumar, Jiewei Jiang).

## Abstract

Chromosomal translocations leading to the fusion of tropomyosin receptor kinases (TRK) with diverse partner proteins have been identified as oncogenic drivers in many adult and pediatric cancers. While first-generation TRK kinase inhibitors, such as entrectinib and larotrectinib, have shown positive responses in TRK fusion-positive cancers, resistance mutations against these inhibitors in the kinase domain limit their efficacy. Second-generation inhibitors are in clinical evaluation, highlighting a need for novel therapeutic modalities to achieve durable suppression of the oncogenic activity of TRK fusions. Here, we developed heterobifunctional small molecule degraders (PROTACs) to achieve targeted degradation of TRK fusions. By conjugating entrectinib to thalidomide, we identified JWJ-01-378 as a potent and selective CRBN-recruiting degrader of the TPM3-TRKA fusion. JWJ-01-378 induced TPM3-TRKA degradation through the ubiquitin-proteasome system and proteomics analysis confirmed the acute selectivity of JWJ-01-378 for achieving TPM3-TRKA degradation with minimal off-target effects. While JWJ-01-378 was also able to degrade wild-type TRK, it was unable to degrade TRK inhibitor resistant mutants and ALK fusions. Importantly, TPM3-TRKA degradation by JWJ-01-378 suppressed downstream signaling and reduced cancer cell viability, with improved responses compared to heterobifunctional control compounds that cannot degrade TPM3-TRKA. Together, our study expands the toolbox of compounds for evaluating targeted degradation of TRK fusions in cancer.

**Graphical Abstract:** 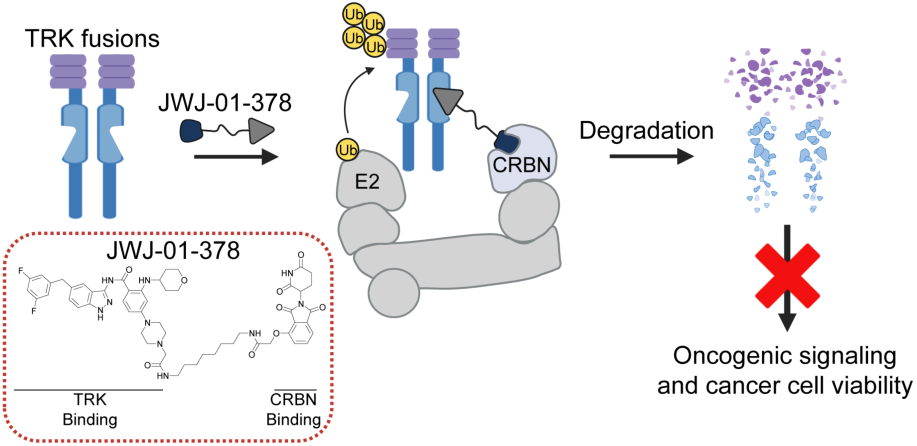

JWJ-01-378 recruits cereblon (CRBN) to induce potent and selective degradation of oncogenic TRK fusions, leading to a collapse in downstream signaling and loss of cancer cell viability. Graphical abstract was created using Biorender.com.

## Introduction

The neurotrophic receptor tyrosine kinase (*NTRK*) gene family, which is comprised of *NTRK1*, *NTRK2*, and *NTRK3*, encodes the tropomyosin receptor kinase proteins TRKA, TRKB, and TRKC, respectively^1^. During normal development, these receptor tyrosine kinases are predominantly expressed in neuronal tissues and activated in response to ligands including NGF, BDNF, NT-3 and NT-4. Ligand binding to their extracellular domains induces receptor dimerization, autophosphorylation of tyrosine kinase domain, and activation of downstream signaling including MAPK, PI3K-AKT, mTOR, and PLCγ pathways that are essential for neuronal development, survival, and differentiation^1, 2^. Alterations in *NTRK* genes, which predominately occur through chromosomal translocations and less commonly include point mutations and deletions, have been identified as driver events in several cancers^3–6^. *NTRK* fusions are recurrent events (> 80%) in rare cancers, such as secretory breast cancer and infantile fibrosarcoma, while less frequent (< 25%) in pediatric and the adult cancers, such as high-grade glioma, melanoma, colorectal cancer, and lung cancer^6^. *In vivo* expression of TRK fusions in mice is sufficient to induce tumor formation, further highlighting their role as strong oncogenic drivers^7, 8^.

Most commonly, these chromosomal translocations involve fusion of the 3’ region of an *NTRK* gene, which encodes the functional tyrosine kinase domain, to the 5’ region of another gene partner^9, 10^. Many of these 5’ fusion partners contribute oligomerization domains that induce ligand-independent receptor dimerization, leading to constitutive activation of the TRK fusions^6,10^. TRK fusions then coordinate aberrant downstream signaling and promote uncontrolled cell proliferation and survival, which are hallmark features of cancer^11^. As such, targeting TRK fusions has emerged as a tractable drug discovery approach, and several TRK inhibitors have advanced into clinical trials.

TRK-targeting compounds include broad-spectrum tyrosine kinase inhibitors such as entrectinib, lestaurtinib, and crizotinib, which demonstrate activity against multiple kinases including TRK, ALK, and ROS1^12–15^. Inhibitors with improved specificity towards TRK proteins have also been developed such as larotrectinib and BPI-28592^16, 17^. Notably, entrectinib and larotrectinib received FDA approval in 2018 and 2019, respectively, for the treatment of cancers harboring TRK mutations and TRK fusions^18, 19^. Both entrectinib and larotrectinib have shown high overall response rates in TRK fusion-positive cancers^20–22^, but the emergence of drug resistance to these inhibitors poses significant clinical challenges^6, 23, 24^. Common resistance mechanisms involve on-target mutations in the TRK kinase domain, including the gatekeeper residue, solvent front and activation loop. Second-generation TRK inhibitors designed to overcome these resistance mutations including selitrectinib, repotrectinib, and ONO-5390556 are currently under clinical evaluation^25–28^. However, on-target mutations including secondary mutations in TRK confer resistance to these inhibitors^29^. Therefore, continued advancement of novel therapeutic approaches is necessary to target TRK fusions in cancer.

Rather than inhibiting protein activity, small molecule degraders such as proteolysis-targeting chimeras (PROTACs) are a powerful chemical strategy to target proteins for degradation through the ubiquitin-proteasome system^30, 31^. PROTACs are heterobifunctional molecules that induce proximity between a target protein and an E3 ubiquitin ligase, leading to ubiquitination and subsequent proteasomal degradation. PROTACs offer several advantages compared to small molecule inhibitors, including their catalytic nature and ability to achieve rapid and sustained elimination of all protein functions including non-enzymatic roles^30^. We^32, 33^ and others^34, 35^ have developed PROTACs targeting kinases, demonstrating their improved activity in dampening aberrant downstream signaling and biological responses compared to kinase inhibitors. Given the oncogenic role of TRK fusions in many pediatric and adult cancers and the challenges associated with resistance mutations limiting the effectiveness of TRK inhibitors, we aimed to develop PROTACs for targeting TRK fusions for degradation.

To study TRK fusion protein degradation, we focused on the TPM3-TRKA fusion and synthesized PROTACs by conjugating entrectinib to thalidomide to recruit the E3 ubiquitin ligase cereblon (CRBN). We identified several compounds as effective degraders, with JWJ-01-378 nominated as the most potent degrader of the TPM3-TRKA fusion, with activity comparable to a previously described TRK PROTAC (CG-428)^36^. JWJ-01-378 required the ubiquitin-proteasome system for achieving degradation and reduced viability of colorectal cancer cells, with enhanced effects compared to heterobifunctional control compounds that cannot degrade TRK. Proteomics analyses confirmed TPM3-TRKA degradation with minimal off-target and secondary responses at an acute time-point. In addition, we observed that JWJ-01-378 and CG-428 degraded wild-type (WT) TRK and were unable to degrade TRK inhibitor resistant mutants or ALK fusions. Together, we developed an expanded toolbox of compounds for the interrogation of TRK fusion proteins in cancer.

## Results and Discussion

### Synthesis of entrectinib-based PROTACs for targeted degradation of TRK fusions

In this study, we aimed to develop TRK PROTACs to evaluate targeted degradation of TRK fusions. As the TRK-binding moiety, we selected entrectinib, which is a potent FDA-approved pan-TRK inhibitor for treating TRK-dependent tumors^37^. In addition, its synthetic simplicity, the commercial availability of an intermediate, and the lack of requirement for chiral separation, made it an attractive starting point for the PROTAC development. Entrectinib binds to the active conformation of TRK kinase by occupying ATP-binding sites in the kinase domain to prevent phosphorylation of its downstream substrates. To functionalize entrectinib to a CRBN-recruiting PROTAC, we evaluated the reported co-crystal structure in complex with TRKA kinase domain (PDB ID: 5KVT) and decided to use solvent exposed *N*-methyl piperazine as a convenient anchor point (**Fig. 1A and B**).

**Figure 1.**
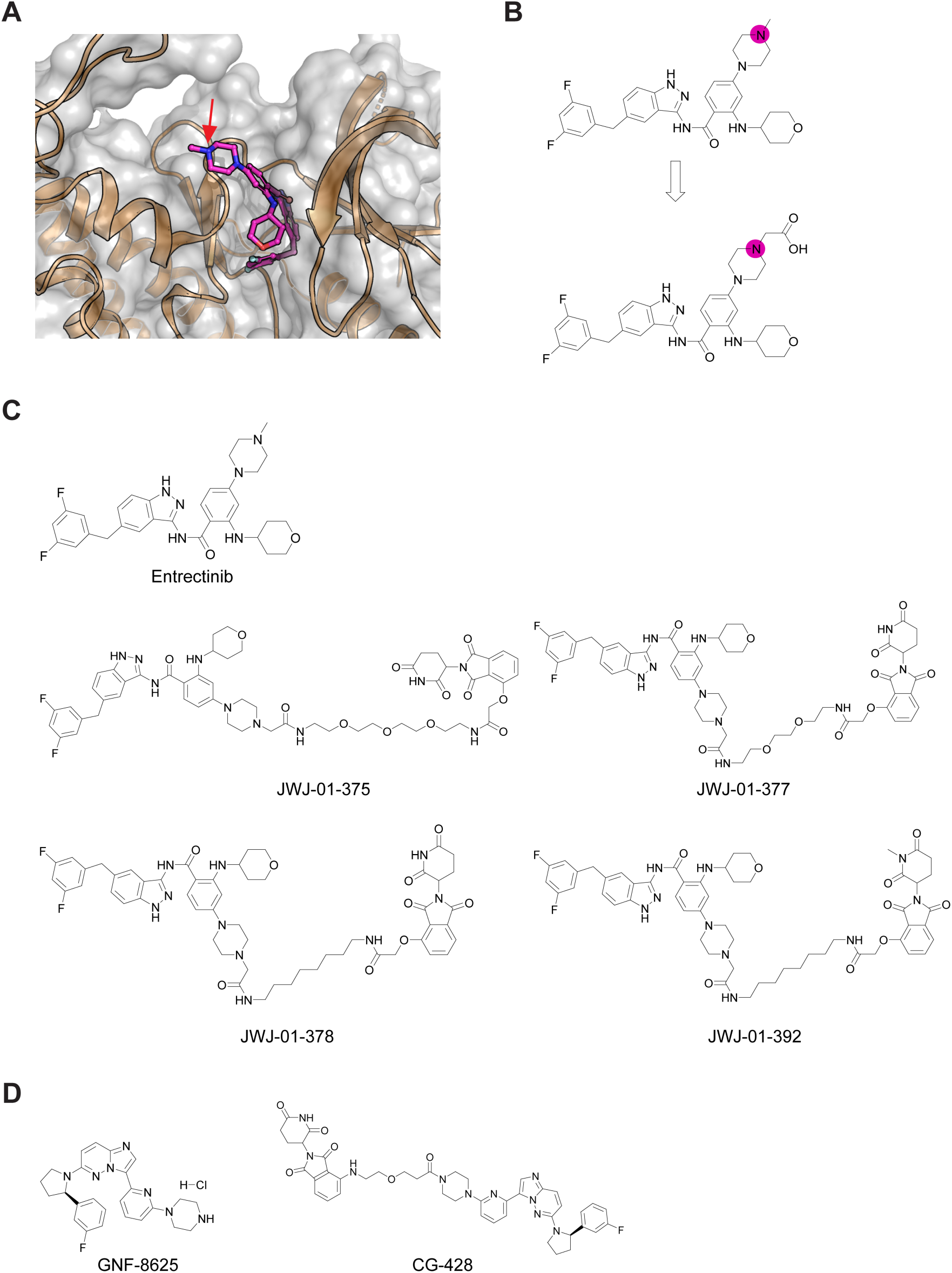
Design and structures of entrectinib-derived PROTACs. (A) The co-crystal structure of entrectinib with TRKA kinase domain (PDB ID: 5KVT), in which the *N*-methyl piperazine is solvent accessible, as indicated by the red arrow. (B) The proposed modification on the piperazine ring for the linker introduction. (C-D) Chemical structures of the indicated compounds.

As depicted in **Scheme 1**, the key carboxylate intermediate was generated within ten steps. Briefly, methyl 2-amino-4-bromobenzoate underwent a reductive amination to introduce the tetrahydropyran ring. A Buchwald-Hartwig coupling installed 1-Boc protected piperazine at the para position of the aryl ring. After the hydrolysis, the secondary aniline was protected by trifluoroacetic anhydride, followed by the oxalyl chloride-mediated amidation with the commercially available indazole moiety. The trifluoroacetic and Boc protecting groups were removed by TEA and TFA, respectively. The displacement between the piperazine and *tert*-butyl bromoacetate and sequential TFA treatment yielded the target carboxylate.

**Scheme 1.**
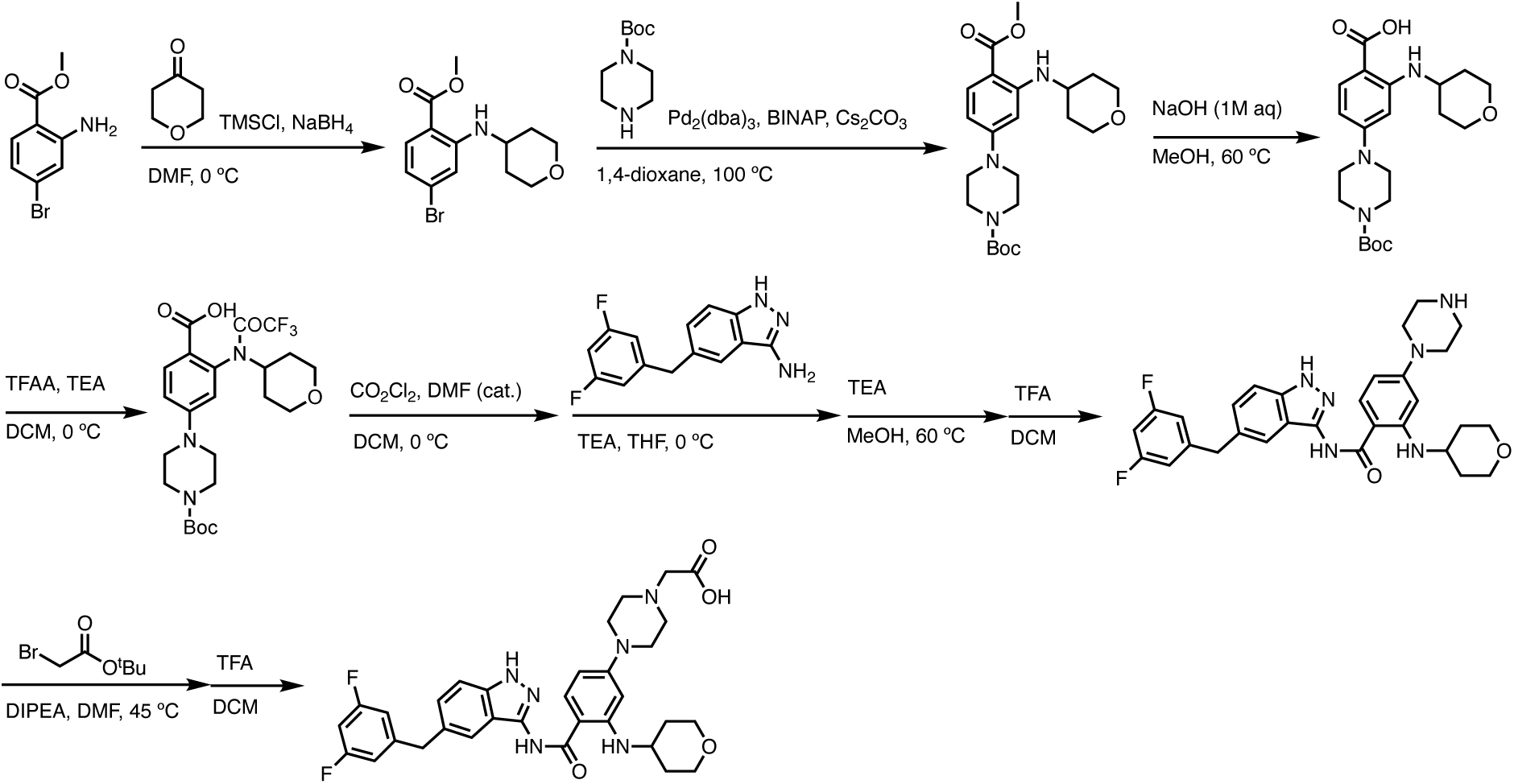
Synthetic route of the key carboxylate intermediate.

Meanwhile as shown in **Scheme 2**, 4 hydroxy thalidomide was treated with *tert*-butyl bromoacetate at the presence of potassium carbonate to selectively alkylate at phenolic site. After the TFA treatment, the carboxylate was coupled with various N-Boc protected linkers. Another Boc deprotection yielded the terminal amine to be integrated with the acid and give the desired PROTAC molecules. Together, we synthesized several entrectinib-derived PROTACs for evaluation (**Fig. 1C and S1A, B, C, and D**).

**Scheme 2.**
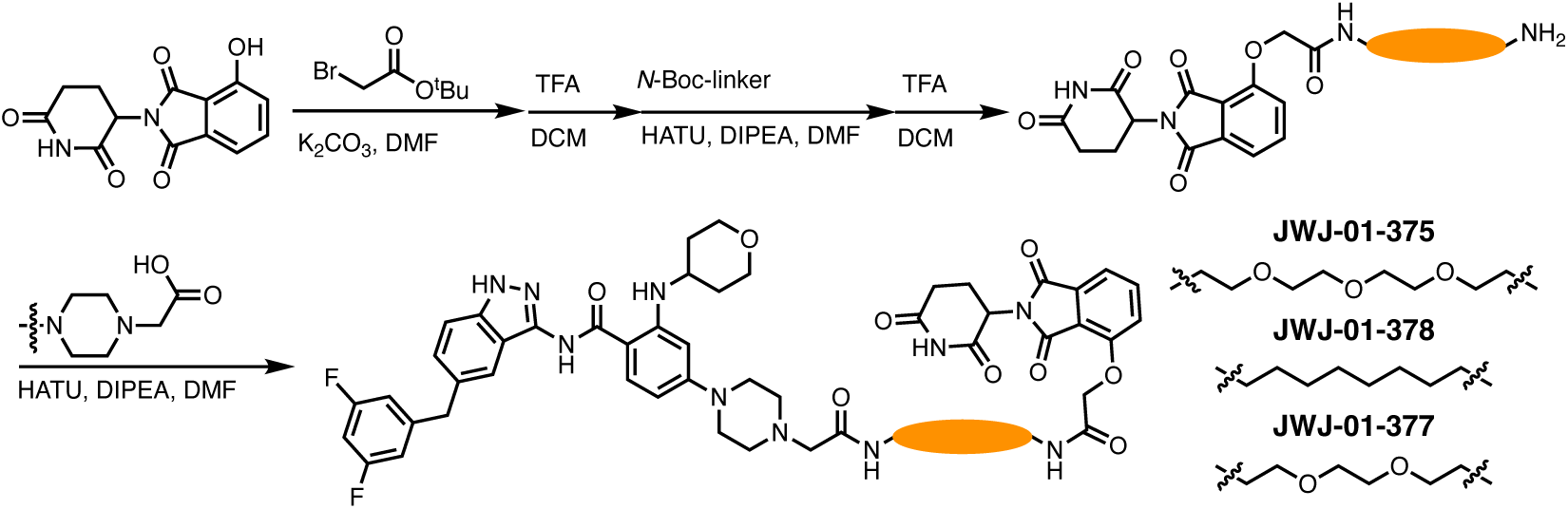
Synthetic route of the designed PROTACs.

### Entrectinib-derived PROTACs induce rapid and potent degradation of TRK fusion

To evaluate the activity of our entrectinib-derived PROTACs, we utilized KM12 cells, a colorectal cancer cell line that harbors an endogenous TPM3-TRKA fusion. This fusion arises from an intrachromosomal rearrangement at 1q22-23 between the N-terminal domain of TPM3 (codons 1-221) and the C-terminal portion of TRKA (codons 419-end) that encodes the transmembrane and intracellular tyrosine kinase domain^38^. As a comparison to our entrectinib-derived PROTACs, we evaluated Cullgen’s TRK-targeting PROTAC (CG-428)^36^, which is composed of GNF-8625 as the TRK-targeting ligand and pomalidomide as the CRBN-recruiting ligand (**Fig. 1D**).

Treatment of KM12 cells with JWJ-01-375 (3-PEG linker), JWJ-01-377 (2-PEG linker), and JWJ-01-378 (8-carbon linker) led to rapid and pronounced degradation of TPM3-TRKA, leading to suppression of phosphorylated ERK (pERK) levels, a pathway that is aberrantly activated by TRK fusions^39^ (**Fig. 2A and B**). Notably, of the PROTACs tested, JWJ-01-378 achieved the most potent degradation at doses as low as 10 nM and was prioritized for further evaluation. As expected, CG-428 and entrectinib decreased pERK levels, and degradation of TPM3-TRKA was observed only upon treatment with CG-428 (**Fig. 2A and B**).To understand and evaluate JWJ-01-378 activity and biological effects, we synthesized a heterobifunctional control compound (JWJ-01-392) consisting of an N-methylated glutarimide analog that disables CRBN binding^40^ (**Fig. 1C**). JWJ-01-392 failed to induce TPM3-TRKA degradation, and did not suppress pERK to the levels of JWJ-01-378 (**Fig. 2C and D**).

**Figure 2.**
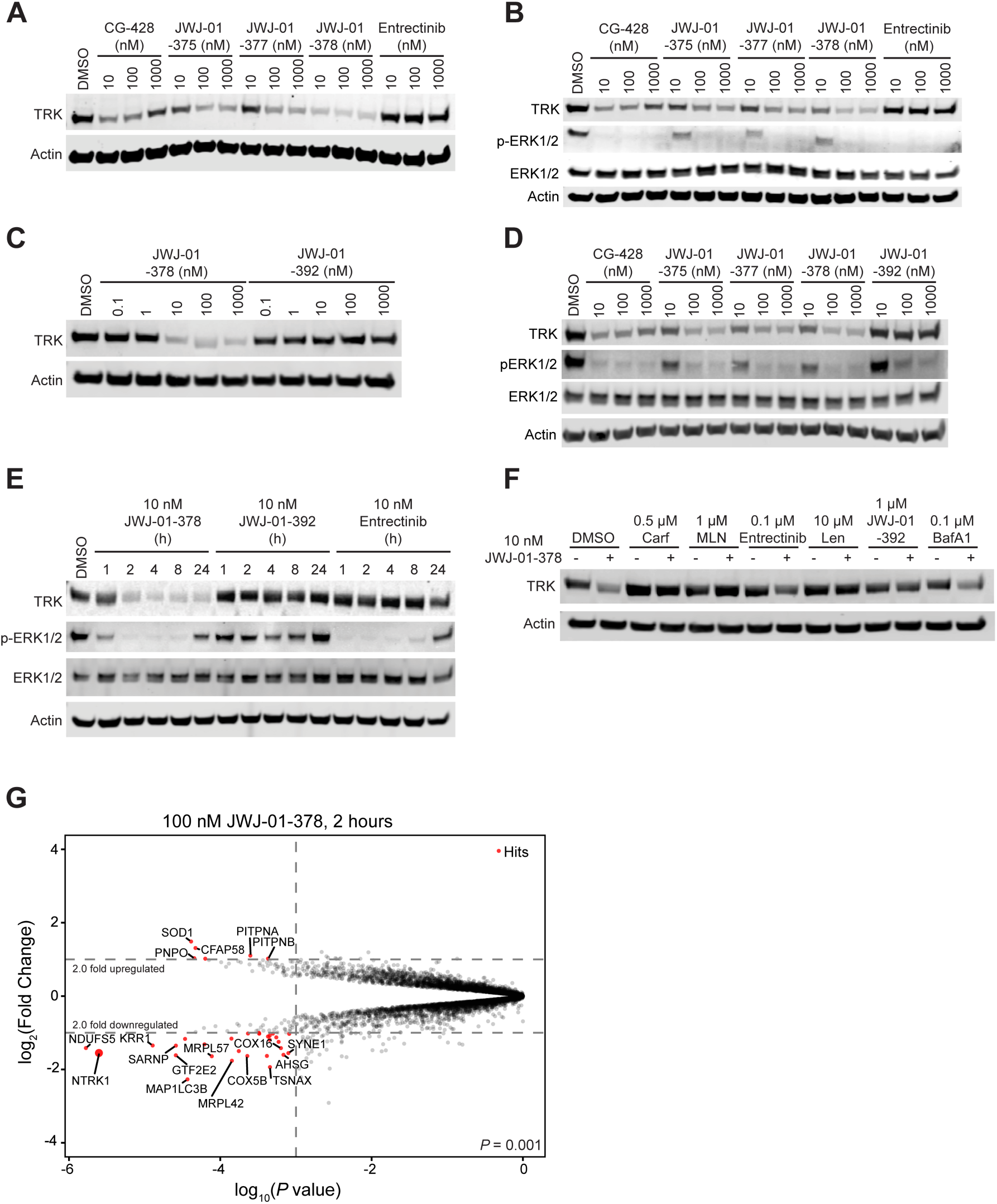
JWJ-01-378 induces potent degradation of TPM3-TRKA. (A) Immunoblot analysis of KM12 cells treated with DMSO, CG-428, JWJ-01-375, JWJ-01-377, JWJ-01-378, and entrectinib at the indicated doses for 2 hours. (B) Immunoblot analysis of KM12 cells treated with DMSO, CG-428, JWJ-01-375, JWJ-01-377, JWJ-01-378, and entrectinib at the indicated doses for 6 hours. (C) Immunoblot analysis of KM12 cells treated with DMSO, JWJ-01-378 and JWJ-01-392 at the indicated doses for 2 hours. (D) Immunoblot analysis of KM12 cells treated with DMSO, CG-428, JWJ-01-375, JWJ-01-377, JWJ-01-378, and JWJ-01-392 at the indicated doses for 6 hours. (E) Immunoblot analysis of KM12 cells treated with DMSO, 10 nM JWJ-01-378, 10 nM JWJ-01-392 and 10 nM entrectinib for the indicated time-course. (F) Immunoblot analysis of KM12 cells pretreated with 0.5 µM of carfilzomib (Carf), 1 µM MLN4924 (MLN), 0.1 µM entrectinib, 10 µM lenalidomide (Len), 1 µM JWJ-01-392, or 0.1 µM bafilomycin A1 (BafA1) for 2 hours followed by DMSO or 10 nM JWJ-01-378 treatment for 4 hours. Data in (A-F) are representative of *n* = 3 independent experiments. (G) Protein abundance after KM12 cells treated with 100 nM of JWJ-01-378 for 2 hours compared to DMSO. Volcano plots depict fold change abundance relative to DMSO versus *P* value derived from a moderated *t*-test. Data are from *n* = 3 independent biological samples.

We next performed time-courses to understand the kinetics of TPM3-TRKA degradation and impacts on downstream pERK signaling. JWJ-01-378 treatment diminished downstream TPM3-TRKA protein levels and abolished pERK signaling, with substantially improved suppression of pERK compared to JWJ-01-392 (**Fig. 2E**). Entrectinib treatment also abolished signaling similar to JWJ-01-378, albeit with slightly faster kinetics, which may be due to improved cell permeability. Interestingly, our time-course analysis revealed partial recovery of pERK levels by 24 hours of treatment with the TRK PROTACs or inhibitors, suggesting the emergence of compensatory signaling or cellular rewiring to reactivate the pathway.

Following confirmation of the disruption of downstream signaling, we confirmed the mechanism-of-action of JWJ-01-378. Pretreatment with proteasomal inhibitor carfilzomib or the NEDD8 inhibitor MLN4924 successfully rescued TPM3-TRKA degradation upon JWJ-01-378 treatment, confirming its dependence on the ubiquitin-proteasome system (**Fig. 2F**). In addition, pretreatment with lenalidomide to saturate CRBN binding or pretreatment with entrectinib or JWJ-01-392 to saturate TRK-kinase domain binding also rescued the degradation of TPM3-TRKA by JWJ-01-378 (**Fig. 2F**). By contrast, inhibition of autophagy with bafilomycin A1 failed to rescue TPM3-TRKA loss upon JWJ-01-378 treatment, further validating that JWJ-01-378 induces TRK degradation through the proteasome (**Fig. 2F**). Together these findings demonstrate that JWJ-01-378 efficiently induces rapid degradation of the TPM3-TRKA fusion protein within two hours, leading to impairment of downstream signaling.

To evaluate the proteome-wide specificity of JWJ-01-378, we performed data-independent acquisition (DIA) proteomic profiling of KM12 cells treated with 100 nM of JWJ-01-378 for two hours (**Fig. 2G**). JWJ-01-378 led to significant TRK fusion degradation after two hours of treatment, with minimal decreases in other proteins. Proteins that displayed alterations included NDUFS5 (NADH dehydrogenase-Complex I), KRR1 (small subunit processome component homolog), SARNP (SAP Domain Containing Ribonucleoprotein), COX16 (mitochondria cytochrome C oxidase assembly), MAP1LC3B (the microtubule-associated protein 1 light chain 3β) and GTF2E2 (general transcription factor 2E2), which may have resulted as a consequence of TPM3-TRKA fusion degradation and rapid changes in downstream signaling that are observed as early as one hour after treatment. Additionally, JWJ-01-378 did not alter the levels of known kinase off-targets of entrectinib, such as ROS1, ALK, JAK, PTK2, SYK, LCK, ABL, KIT, and FGFR. Together these data confirm that JWJ-01-378 is a fast-acting, selective, and potent degrader of TPM3-TRKA.

### Entrectinib-derived PROTACs degrade wild-type TRK but not TRK inhibitor resistance mutants or EML4-ALK

As entrectinib has been shown to inhibit both WT TRK and TRK fusions^41^, we sought to evaluate WT TRK degradation in human erythroleukemia (HEL) cells, which endogenously express WT TRKA, and NIH/3T3 cells ectopically expressing WT TRKA. Treatment with JWJ-01-378 and CG-428 resulted in a significant reduction in WT TRKA protein levels, whereas JWJ-01-392 had no effect (**Fig. 3A and B**). We next evaluated whether these PROTACs were able to degrade TRK-inhibitor resistance mutants. The G595R point mutation confers resistance to entrectinib^42^, which we expressed in NIH/3T3 cells and compared the degradation to TPM3-TRKA. While we observed degradation of TPM3-TRKA, JWJ-01-378 and CG-428 failed to induce degradation of TPM3-TRKA-G595R (**Fig. 3C and D**). In addition, a reported off-target of entrectinib is ALK^13^, and we evaluated whether entrectinib-derived PROTACs were able to degrade ALK fusions in NIH/3T3 cells ectopically expressing EML4-ALK or a drug-refractory mutant EML4-ALK-G1202R^43^. At the doses and time-points assessed, we did not observe degradation of EML4-ALK or EML4-ALK-G1202R upon treatment with JWJ-01-378 or CG-428 (**Fig. 3E, F**, and **G**). Together, these findings indicate JWJ-01-378 and CG-428 degrade WT TRK and TPM3-TRKA and are unable to degrade TPM3-TRKA-G595R, EML4-ALK, or EML4-ALK-G1202R.

**Figure 3.**
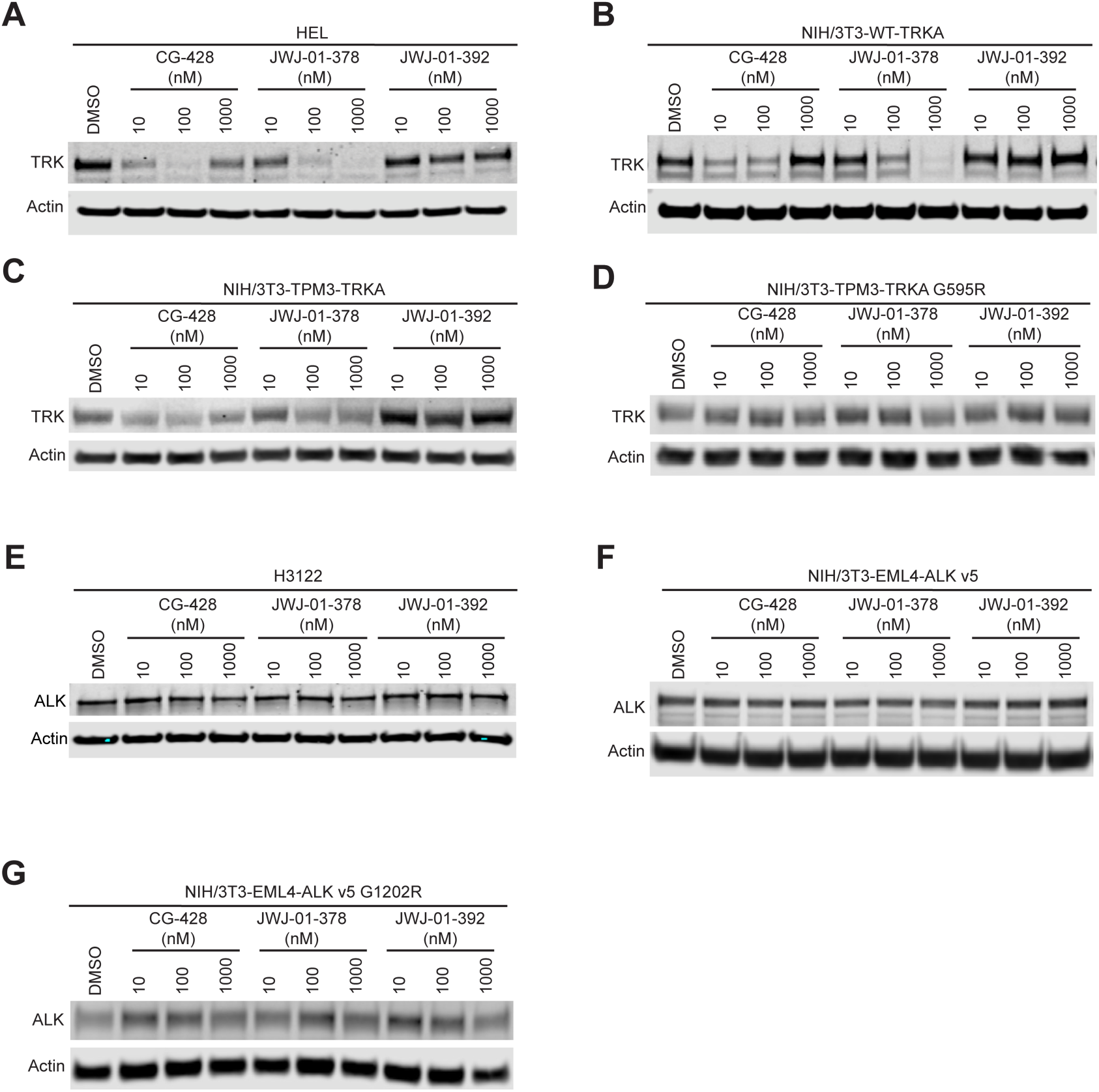
TRK PROTACs degrade WT TRKA but not TPM3-TRKA-G595R, EML4-ALK, or EML4-ALK-G1202R. (A) Immunoblot analysis of HEL cells treated with DMSO, CG-428, JWJ-01-378, and JWJ-01-392 at the indicated doses for 6 hours. (B) Immunoblot analysis of NIH/3T3 cells expressing WT-TRKA treated with DMSO, CG-428, JWJ-01-378, and JWJ-01-392 at the indicated doses for 6 hours. (C) Immunoblot analysis of NIH/3T3 cells expressing TPM3-TRKA treated with DMSO, CG-428, JWJ-01-378, and JWJ-01-392 at the indicated doses for 6 hours. (D) Immunoblot analysis of NIH/3T3 cells expressing TPM3-TRKA-G595R treated with DMSO, CG-428, JWJ-01-378, and JWJ-01-392 at the indicated doses for 6 hours. (E) Immunoblot analysis of H3122 cells treated with DMSO, CG-428, JWJ-01-378, and JWJ-01-392 at the indicated doses for 6 hours. (F) Immunoblot analysis of NIH/3T3 cells expressing EML4-ALK treated with DMSO, CG-428, JWJ-01-378, and JWJ-01-392 at the indicated doses for 6 hours. (G) Immunoblot analysis of NIH/3T3 cells expressing EML4-ALK-G1202R treated with DMSO, CG-428, JWJ-01-378, and JWJ-01-392 at the indicated doses for 6 hours. Data in (A-G) are representative of *n* = 3 independent experiments.

### JWJ-01-378 diminishes viability of TRK fusion-positive cells

To assess the biological consequences of PROTAC-mediated degradation of TPM3-TRKA, we evaluated the antiproliferative effects of our compounds in 2D-adherent monolayer and 3D-spheroid cultures. In particular, 3D-spheroids are valuable models for providing predictive insights into *in vivo* efficacy^8, 32^. KM12 cells were treated with a panel of TRK inhibitors (entrectinib and GNF-8625), PROTACs (JWJ-01-375, JWJ-01-377, JWJ-01-378, and CG-428), and heterobifunctional controls (JWJ-01-392) (**Fig. 4A, B, C, and D and Table S1**). In line with the observed degradation effects, JWJ-01-378 demonstrated the most potent antiproliferative activity in both 2D-adherent monolayer and 3D-spheroid cultures, compared to JWJ-01-375 and JWJ-01-377, with a substantial shift in IC_50_ compared to JWJ-01-392. However, JWJ-01-378 was less effective than entrectinib, GNF-8625, and CG-428, which may be due to higher cell permeability, contributions from polypharmacology, efflux of JWJ-01-378, or superior ability of CG-428 to sustain degradation over 120 hours. Having observed that TRK PROTACs degrade WT TRK (**Fig. 3A and B**), we also tested whether TRK PROTACs or inhibitors reduced the viability of HEL cells, which express WT TRK and do not harbor an TRK fusion, and did not observe an antiproliferative effect in this model (**Fig. S2A and B**).

**Figure 4.**
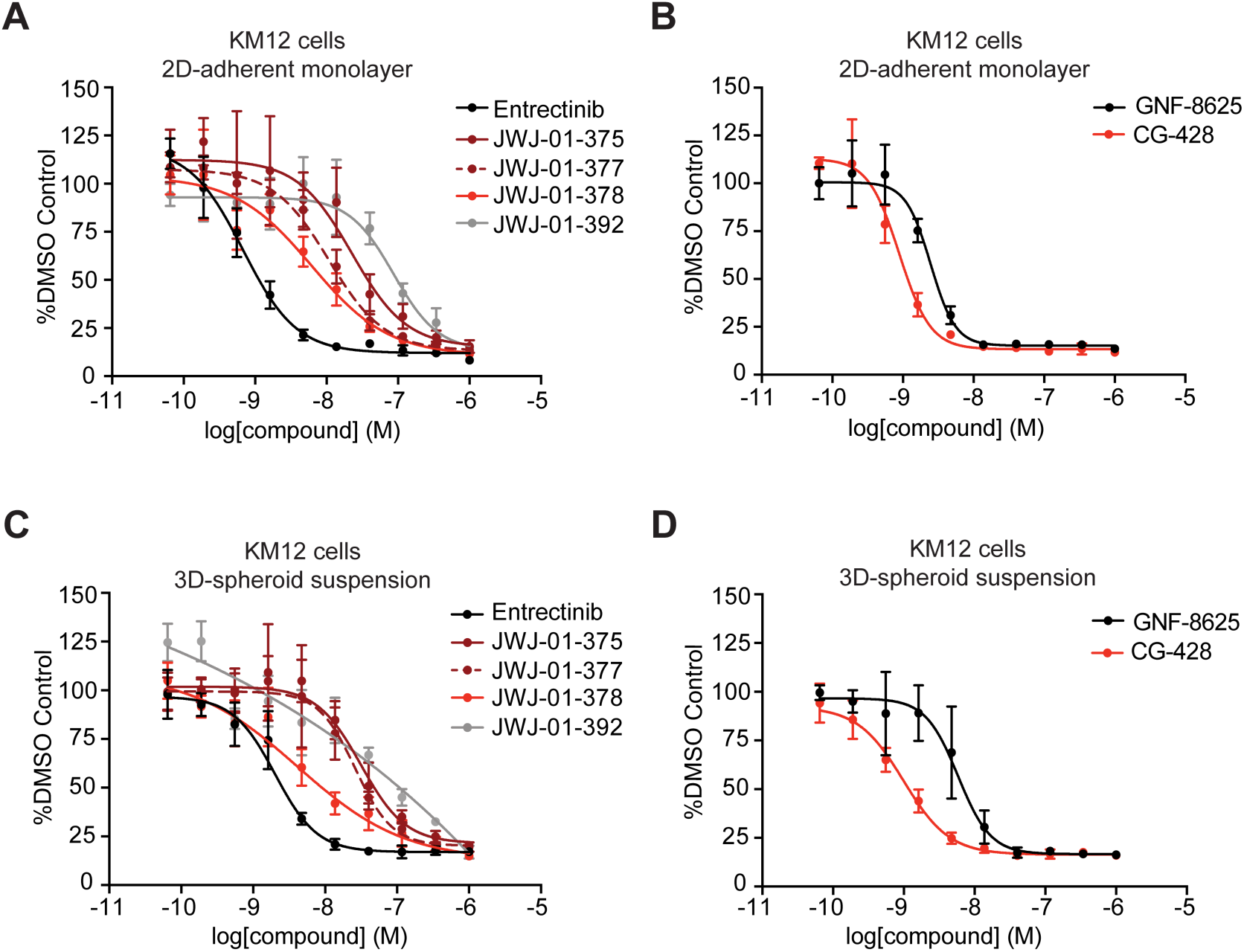
JWJ-01-378 diminishes proliferation of TPM3-TRKA cells. (A-B) DMSO-normalized antiproliferation of KM12 cells cultured as 2D-adherent monolayers and treated with indicated compounds for 120 hours. (C-D) DMSO-normalized antiproliferation of KM12 cells cultured as ultra-low adherent 3D spheroid suspension conditions and treated with indicated compounds for 120 hours. Data in (A-D) are presented as mean ± s.d. of *n* = 4 biologically independent samples and are representative of *n* = 3 independent experiments.

Drug efflux has emerged as a common mechanism of diminishing PROTAC activity^44, 45^. To test whether ABCB1 transporters were contributing to these differences, we co-treated entrectinib, JWJ-01-378 or CG-428 with the ABCB1-efflux pump inhibitors tariquidar (**Fig. S3A**) and zosuquidar (**Fig. S3B**). We also evaluated whether a second treatment of JWJ-01-378, CG-428 or entrectinib at 72 hours improved responses (**Fig. S3C**). Co-treatment with tariquidar or zosuquidar or a second treatment of the compounds did not substantially improve the antiproliferative effects of the compounds. Together, these data suggest that JWJ-01-378 diminishes the proliferation of TPM3-TRKA fusion-positive cells, with improved effects compared to JWJ-01-392, its matched heterobifunctional control compound.

## Conclusions

TRK fusions are oncogenic drivers of several adult and pediatric cancers. First-generation and second-generation TRK inhibitors have demonstrated efficacy in clinical trials in patients with TRK-driven malignancies including SCOUT (NCT02637687) and NAVIGATE (NCT02576431)^6, 46^. However, these inhibitors have off-target effects and the development of acquired resistance limits their long-term clinical utility. Targeted protein degradation is a promising approach to circumvent these limitations by promoting the complete removal of wild-type and mutant proteins^47–50^. In this study, we aimed to develop PROTACs to target TRK fusion proteins for degradation. Using structure-guided design, we conjugated entrectinib to thalidomide and nominated JWJ-01-378 as a potent TRK-targeting PROTAC. JWJ-01-378 induces the degradation of WT TRK and TRK fusions within 2 hours of treatment at a 10 nM dose, diminishes levels of downstream pERK signaling, and requires the ubiquitin-proteasome system for degradation. Proteomics analysis confirmed the specificity of TPM3-TRKA degradation at two hours.

We also compared JWJ-01-378 to inhibitors (entrectinib and GNF-8625), CG-428, and a heterobifunctional control compound (JWJ-01-392). While JWJ-01-378 achieved substantial TPM3-TRKA degradation, comparable to CG-428, it did not diminish the viability of KM12 cells to levels observed with entrectinib, GNF-8625, or CG-428. These differences in the potency were not due to ABCB1-mediated efflux and were not improved with a second administration of TRK PROTACs. The improved antiproliferative effects upon entrectinib and CG-428 treatment may be due to differential impacts on downstream pathways or off-target effects^51^ that are spared by JWJ-01-378. Importantly, JWJ-01-378 had improved antiproliferative activity in 2D-monolayer and 3D-spheroids compared to JWJ-01-392, supporting the enhanced degradation-dependent effects over a matched heterobifunctional control compound. Notably, while JWJ-01-378 and CG-428 also degrade WT TRK in HEL cells, they did not reduce the viability of these cells. This may be due to dependency of these cells on other factors for survival. For example, HEL cells have an oncogenic JAK2 V617F mutation^52, 53^. The degradation of WT TRK presents an interesting opportunity for an application towards chronic pain management, as WT TRK is a known mediator of chronic pain and nociception^54^.

With the compounds developed in this study, there are several future directions to pursue that were not addressed in this work. First, it will be important to determine how TRK fusions degradation impacts downstream signaling networks to nominate potential drug combination strategies. In our time-course experiments with JWJ-01-378 and entrectinib, we observed reactivation of pERK signaling as early as 24 hours after treatment. Compensatory reactivation of the MAPK pathway is a well-documented mechanism of resistance to TRK inhibitors and may involve upregulation of alternative receptor tyrosine kinases or secondary mutations including BRAF^V600E^, KRAS^G12D^, and amplification of *MET*^42^. Evaluation of combination of strategies with MAPK or PI3K pathway inhibitors to enhance the durability of TRK degradation effects is necessary, which has been previously shown in the context of TRK inhibitors^55^.

Next, it will be important to define the mechanisms of resistance to TRK PROTACs. Although JWJ-01-378 and CG-428 efficiently degraded WT and TPM3-TRKA fusion proteins, they failed to degrade the TPM3-TRKA-G595R resistance mutant. This finding aligns with previous studies showing that the G595R mutation in the kinase domain of TRK confers resistance to TRK inhibitors, such as entrectinib^23^. Furthermore, identifying the spectrum of TRK fusions targeted by JWJ-01-378 due to binding the TRK kinase domain will be important to assess its ability to tackle the EML4-TRK, ETV6-TRK, and SQSTM1-TRK fusions that are observed in many cancers. Interestingly, while ALK is a known off-target of entrectinib, JWJ-01-378 did not degrade EML4-ALK or EML4-ALK-G1202R, suggesting selectivity for TRK over ALK proteins, and the necessity of these evaluations. Finally, continued evaluation of TRK-targeting PROTACs in mouse models of cancer is necessary to understand toxicities and potential as a therapeutic approach in cancer. In conclusion, JWJ-01-378 represents a potent TRK-targeting PROTAC that efficiently reduces TPM3-TRKA fusion protein levels and suppresses cancer cell proliferation. However, its inability to degrade TRK resistance mutants and the activation of compensatory MAPK signaling responses highlight the need for further optimization and combinatorial treatment strategies. These findings provide a strong foundation for the continued development of TRK-targeting PROTACs as therapeutic agents for TRK fusion-driven cancers.

## Author Contributions

SK and JJ contributed equally to this work. SK, MSDP, and JC performed all the biological experiments. JJ performed compound design and synthesis and docking modeling. AG and AJF performed proteomics studies and AG, AJF, and SK analyzed the data. FS and ECH provided resources and analyzed and interpreted the data. FMF and BN conceptualized the study, designed the experiments, analyzed and interpreted the data, acquired funding, and supervised the study. SK and BN wrote the manuscript with input from all authors.

## Data availability

The raw proteomics datasets generated during this study are available at the ProteomeXchange Consortium via the PRIDE partner repository^56, 57^ with the dataset identifier PXD065035.

## Acknowledgements

The authors thank members of the Nabet and Ferguson laboratories for feedback on this study and manuscript. This work was supported by: Hartwell Innovation Fund – Swim Across America (SK), NSF CAREER CHE-2339705 (FMF), NIH/NCI K22 CA258805 (BN), NIH/NCI Cancer Center Support Grant P30 CA015704 (BN), NIH/NCI U01 CA282109 (BN), and NIH/NCATS R43 TR004221 (AJF and BN). This research was supported by the Cellular Imaging Shared Resource, RRID:SCR_022609 of the Fred Hutch/University of Washington/Seattle Children’s Cancer Consortium (P30 CA015704).

## Methods

### Synthetic procedures

#### General methods

Unless otherwise noted, reagents and solvents were obtained from commercial suppliers and were used without further purification. ^11^H NMR spectra were recorded on 500 MHz Bruker Avance III spectrometer or 500 MHz JEOL ECA 500 spectrometer and chemical shifts are reported in parts per million (ppm, δ) downfield from tetramethylsilane (TMS). Coupling constants (J) are reported in Hz. Spin multiplicities are described as s (singlet), br (broad singlet), d (doublet), t (triplet), q (quartet), and m (multiplet). Mass spectra were obtained on a Waters Acquity UPLC/MS. Preparative HPLC was performed on a Waters Sunfire C18 column (19 mm × 50 mm, 5 μM) using a gradient of 15−95% methanol in water containing 0.05% trifluoroacetic acid (TFA) over 22 min (28 min run time) at a flow rate of 20 mL/min. Assayed compounds were isolated and tested as TFA salts. Purities of assayed compounds were in all cases greater than 90%, as determined by reverse-phase HPLC analysis.

**Methyl 4-Bromo-2-((tetrahydro-2*H*-pyran-4-yl)amino)benzoate**

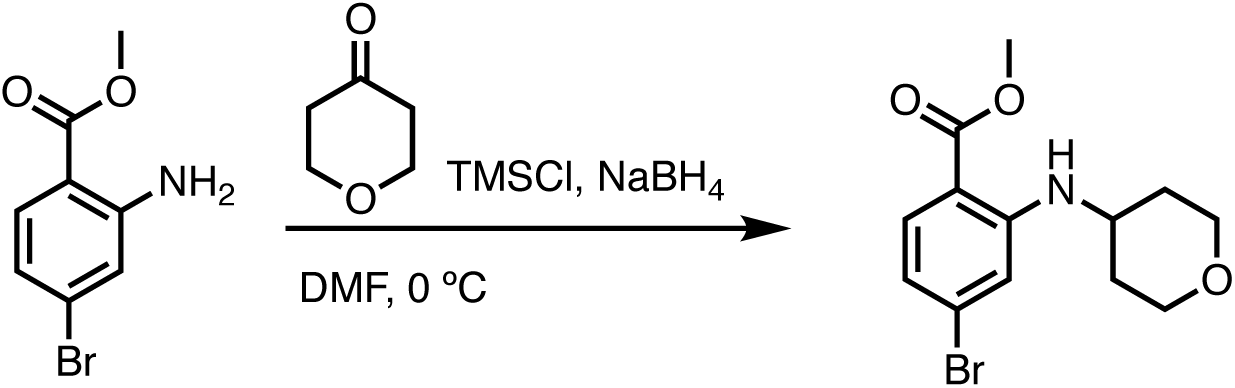

According to a reported literature^58^, a dry 100 mL two-neck round bottom flask with magnetic stirring bar was charged consecutively with methyl 2-amino-4-bromobenzoate (1.0 equiv), tetrahydro-4h-pyran-4-one (1.1 equiv), TMSCl (2.5 equiv), and dry DMF in a nitrogen atmosphere. The reaction mixture was cooled to 0 °C and NaBH_4_ (1.0 equiv) was added in one portion. The reaction mixture was kept stirring at 0 °C until full conversion of the arylamine was detected by

UPLC. The vigorously stirred reaction mixture was treated with saturated NaHCO_3_ solution, followed by EtOAc extraction. The two-phase mixture was kept stirring until the gas evolution had ceased. The phases were separated, and the aqueous layer was further extracted with EtOAc once. The combined organic layers were dried over anhydrous MgSO_4_. The drying agent was removed by filtration, and the solvent was removed under reduced pressure. The crude product was loaded on the silica gel and purified by chromatography, which gave the title compound as pale-yellow solid. ^1^H NMR (500 MHz, DMSO-*d*_6_) δ 7.79 (d, *J* = 7.8 Hz, 1H), 7.70 (dd, *J* = 8.6, 1.1 Hz, 1H), 7.07 (d, *J* = 1.8 Hz, 1H), 6.74 (dt, *J* = 8.6, 1.5 Hz, 1H), 3.83 (dt, *J* = 11.6, 3.7 Hz, 2H), 3.79 (d, *J* = 1.1 Hz, 3H), 3.77–3.69 (m, 1H), 3.53–3.45 (m, 2H), 1.96–1.88 (m, 2H), 1.44–1.34 (m, 2H). MS (ESI) m/z 314.2 (M + H)^+^.

***tert*-Butyl 4-(4-(methoxycarbonyl)-3-((tetrahydro-2*H*-pyran-4-yl)amino)phenyl)piperazine-1-carboxylate**

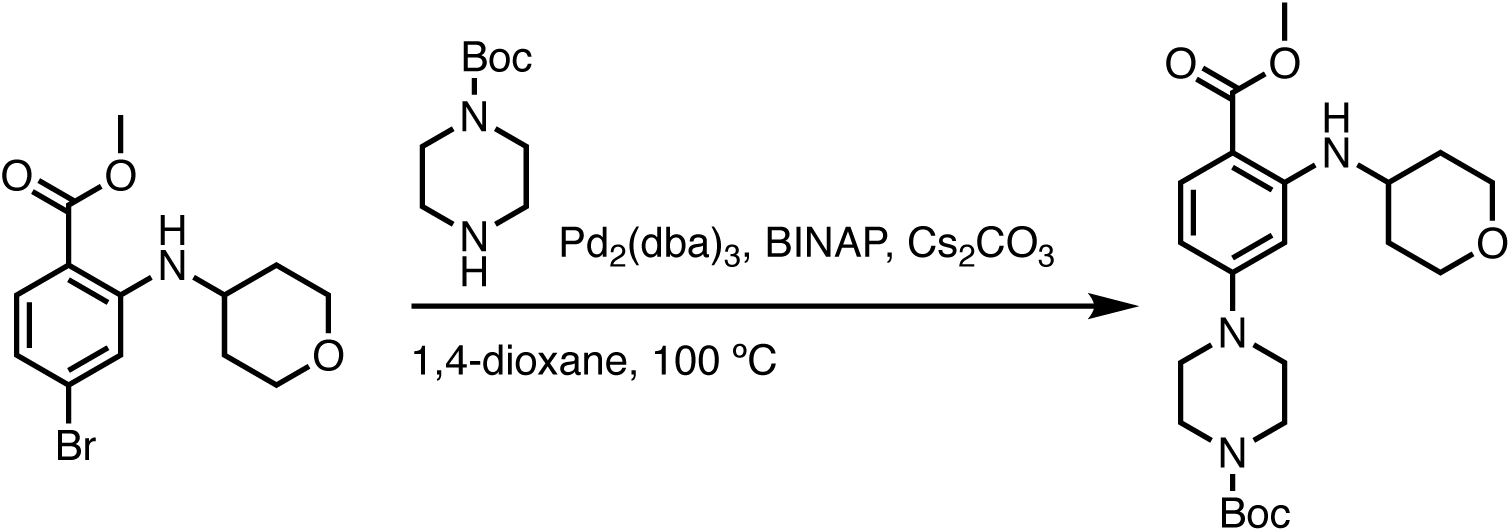

To a two-neck round bottom flask equipped with a nitrogen balloon and a septum, the resulting reductive amination intermediate (1.0 equiv), BINAP (0.2 equiv), Pd_2_(dba)_3_ (0.1 equiv), Cs_2_CO_3_ (3.0 equiv), and *tert*-butyl piperazine-1-carboxylate (1.1 equiv) were charged. Anhydrous 1,4-dixoane was then added and the system was flushed with nitrogen gas three times. The mixture was heated to 100 °C and stirred overnight. Upon complete consumption of reductive amination intermediate monitored by UPLC, 50 mL EtOAc was added and filtered through a pad of Celite. The filtrate was saved and washed with brine twice. The organic layer was dried over anhydrous

MgSO_4_, concentrated under reduced pressure, and purified via flash column chromatography, which provided the title compound as pale-yellow solid. ^1^H NMR (500 MHz, DMSO-*d*_6_) δ 7.76 (d, *J* = 7.7 Hz, 1H), 7.63 (dd, *J* = 9.2, 2.6 Hz, 1H), 6.21 (dd, *J* = 9.3, 2.5 Hz, 1H), 6.08 (d, *J* = 2.4 Hz, 1H), 3.83 (dt, *J* = 11.7, 3.5 Hz, 2H), 3.71 (d, *J* = 2.5 Hz, 4H), 3.53–3.47 (m, 2H), 3.43 (s, 4H), 3.27 (t, *J* = 5.5 Hz, 4H), 1.96 (t, *J* = 12.1 Hz, 2H), 1.42 (d, *J* = 2.6 Hz, 11H). MS (ESI) m/z 420.5 (M + H)^+^.

**4-(4-(*tert*-Butoxycarbonyl)piperazin-1-yl)-2-((tetrahydro-2*H*-pyran-4-yl)amino)benzoic Acid**

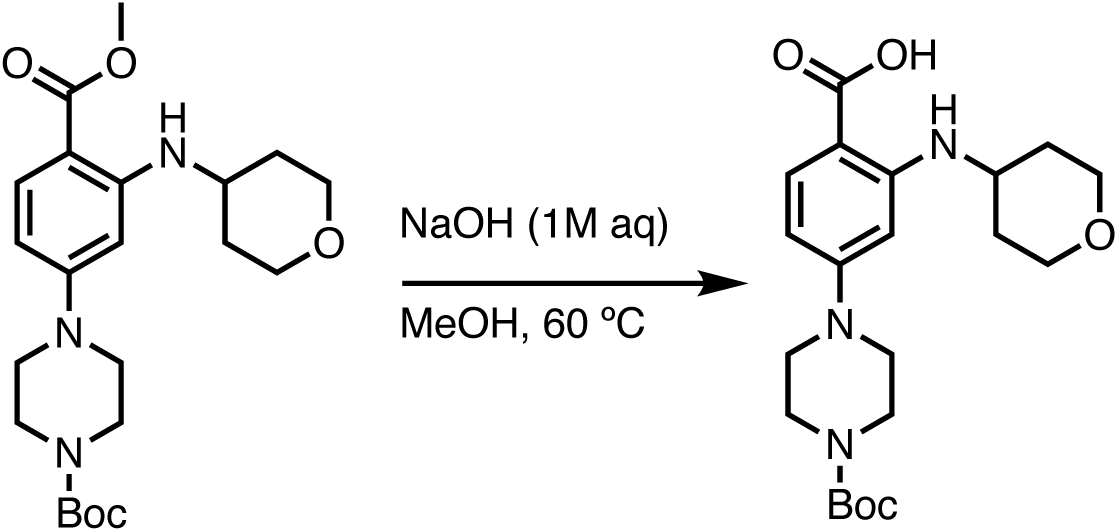

The methyl ester intermediate (1.0 equiv) was dissolved in methanol before the addition of 1M NaOH solution (3.0 equiv). The mixture was stirred for 8 hours at 60 °C, which resulted in a brown clear solution. The solution was then chilled on ice and acidified with 1M HCl solution to pH 2-3, which generated a lot of solid. The cloudy mixture was further stirred for 30 minutes before the filtration to collect the solid. The pale-white solid was dried under reduced pressure to yield the title acid intermediate without further purification. ^1^H NMR (500 MHz, DMSO-*d*_6_) δ 11.93 (s, 1H), 7.93 (d, *J* = 7.6 Hz, 1H), 7.61 (d, *J* = 9.0 Hz, 1H), 6.18 (dd, *J* = 9.1, 2.3 Hz, 1H), 6.07 (d, *J* = 2.4 Hz, 1H), 3.83 (dt, *J* = 11.7, 3.8 Hz, 2H), 3.70 (d, *J* = 11.0 Hz, 1H), 3.49 (td, *J* = 11.2, 2.4 Hz, 2H), 3.43 (d, *J* = 10.6 Hz, 4H), 3.24 (t, *J* = 5.3 Hz, 4H), 1.94 (d, *J* = 12.9 Hz, 2H), 1.41 (s, 9H), 1.35 (ddd, *J* = 13.8, 8.6, 3.9 Hz, 2H). MS (ESI) m/z 406.6 (M + H)^+^.

**4-(4-(*tert*-Butoxycarbonyl)piperazin-1-yl)-2-(2,2,2-trifluoro-*N*-(tetrahydro-2*H*-pyran-4-yl)acetamido)benzoic** Acid

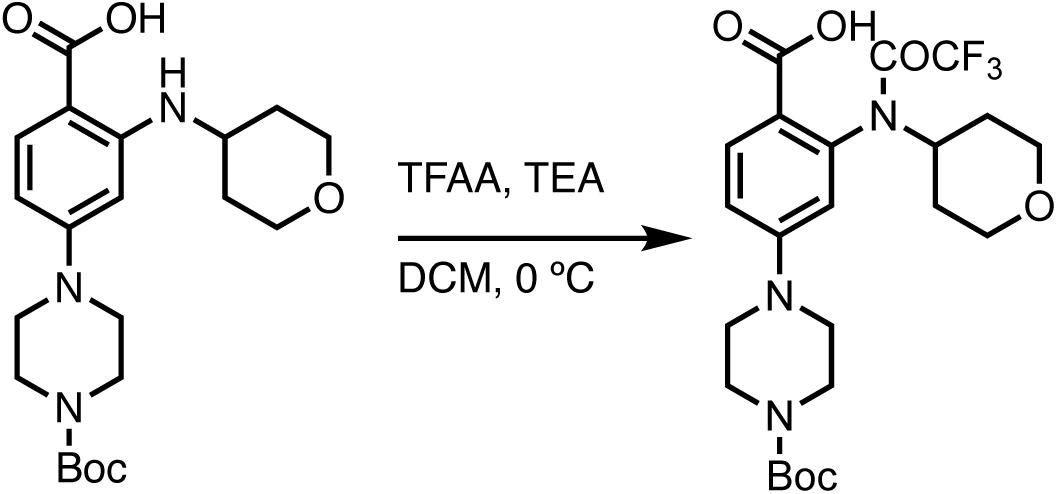

To the acid intermediate (1.0 equiv) solution in DCM, TEA (2.0 equiv) was added. The solution was cooled to 0 °C before the addition of TFAA (1.5 equiv). The reaction was kept stirring at 0 °C until full conversion of the acid was detected by UPLC. The solvent was removed under reduced pressure, which yielded the desired protected intermediate as yellow oil with no further purification. MS (ESI) m/z 502.3 (M + H)^+^.

***tert*-Butyl 4-(4-((5-(3,5-difluorobenzyl)-1*H*-indazol-3-yl)carbamoyl)-3-((tetrahydro-2*H*-pyran-4-yl)amino)phenyl)piperazine-1-carboxylate**

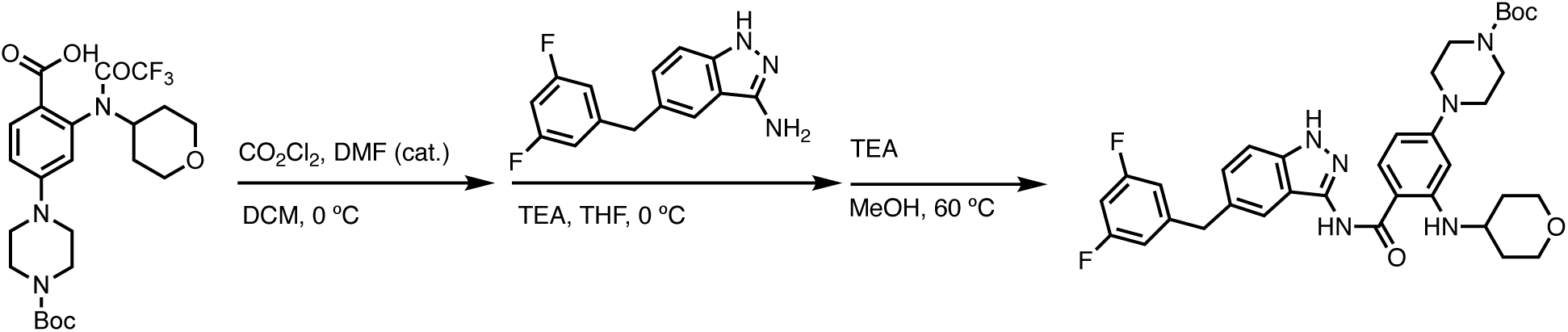

According to a reported literature^19^, to a suspension of acid trifluoroacetate (1.0 equiv) in dry DCM, oxalyl chloride (1.5 equiv) and DMF (1-2 drops, catalytic amount) were added. The mixture was stirred at room temperature for 2 hours then evaporated to dryness. The resulting crude acyl chloride was taken-up with toluene, evaporated, and then dissolved in dry THF at 0 °C. A solution of 5-(3,5-difluoro-benzyl)-1H-indazol-3-ylamine (1.0 equiv) and DIPEA (4.0 equiv) in dry THF was added to the cooled acyl chloride solution. The mixture was stirred at 0 °C for 4 hours then quenched by adding DI H_2_O/EtOAc. The organic phase was washed with a saturated NaHCO_3_ solution, dried over MgSO_4_, and evaporated to dryness. Crude reaction mixture is dissolved in methanol in the presence of triethylamine (3.0 equiv) and stirred at 65 °C for 2 hours. The solvent was removed under reduced pressure and the residue treated with DI H_2_O/EtOAc. Organic layer was dried over anhydrous MgSO_4_ and evaporated to dryness. The crude product was further purified by chromatography over silica gel to generate the title compound as a pale yellow solid. ^1^H NMR (500 MHz, DMSO-*d*_6_) δ 12.66 (s, 1H), 10.12 (s, 1H), 8.29 (d, *J* = 7.6 Hz, 1H), 7.81 (d, *J* = 9.0 Hz, 1H), 7.48 (s, 1H), 7.41 (d, *J* = 8.6 Hz, 1H), 7.25 (dd, *J* = 8.6, 1.6 Hz, 1H), 7.00 (ddd, *J* = 15.5, 7.8, 4.4 Hz, 3H), 6.24 (dd, *J* = 9.0, 2.3 Hz, 1H), 6.15 (d, *J* = 2.4 Hz, 1H), 4.05–4.02 (m, 2H), 3.81 (dt, *J* = 11.8, 3.8 Hz, 2H), 3.68 (d, *J* = 6.6 Hz, 1H), 3.51–3.41 (m, 6H), 3.25 (t, *J* = 5.3 Hz, 4H), 1.94 (d, *J* = 12.8 Hz, 2H), 1.43 (s, 9H), 1.33 (q, *J* = 13.5, 11.6 Hz, 2H). MS (ESI) m/z 647.5 (M + H)^+^.

***tert*-Butyl 2-(4-(4-((5-(3,5-difluorobenzyl)-1*H*-indazol-3-yl)carbamoyl)-3-((tetrahydro-2*H*-pyran-4-yl)amino)phenyl)piperazin-1-yl)acetate**

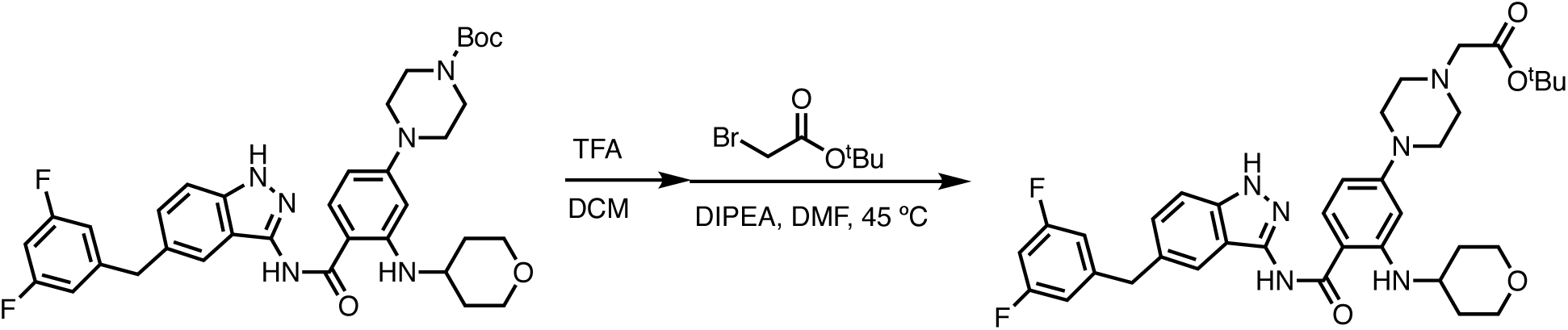

To the Boc-protected intermediate (1.0 equiv) in DCM, TFA (2.0 equiv) was added. The resulting solution was stirred for 1 hour to fully remove the Boc group, indicated by UPLC. The solvent was then removed under reduced pressure before re-dissolving in dry DMF. *tert*-Butyl bromoacetate (1.1 equiv) and DIPEA (1.5 equiv) were added to the solution and stirred for another 1 hour at 45 °C. Upon complete consumption of free piperazine intermediate monitored by UPLC, DI H_2_O/EtOAc were added. The organic layer was dried over anhydrous MgSO_4_, concentrated under reduced pressure, and purified via flash column chromatography, which provided the title compound as pale-yellow solid. ^1^H NMR (500 MHz, DMSO-*d*_6_) δ 12.65 (s, 1H), 10.10 (s, 1H), 8.28 (d, *J* = 7.6 Hz, 1H), 7.79 (d, *J* = 9.0 Hz, 1H), 7.48 (s, 1H), 7.40 (d, *J* = 8.6 Hz, 1H), 7.25 (dd, *J* = 8.6, 1.6 Hz, 1H), 7.03–6.96 (m, 3H), 6.23 (dd, *J* = 9.0, 2.3 Hz, 1H), 6.13 (d, *J* = 2.3 Hz, 1H), 4.03 (s, 2H), 3.81 (dt, *J* = 11.6, 3.9 Hz, 2H), 3.71–3.64 (m, 1H), 3.52–3.46 (m, 2H), 3.27 (t, *J* = 4.9 Hz, 4H), 3.17 (s, 2H), 2.62 (t, *J* = 5.0 Hz, 4H), 1.93 (d, *J* = 12.8 Hz, 2H), 1.43 (s, 9H), 1.33 (td, *J* = 13.6, 9.9 Hz, 2H).

***tert*-Butyl 2-((2-(2,6-dioxopiperidin-3-yl)-1,3-dioxoisoindolin-4-yl)oxy)acetate**

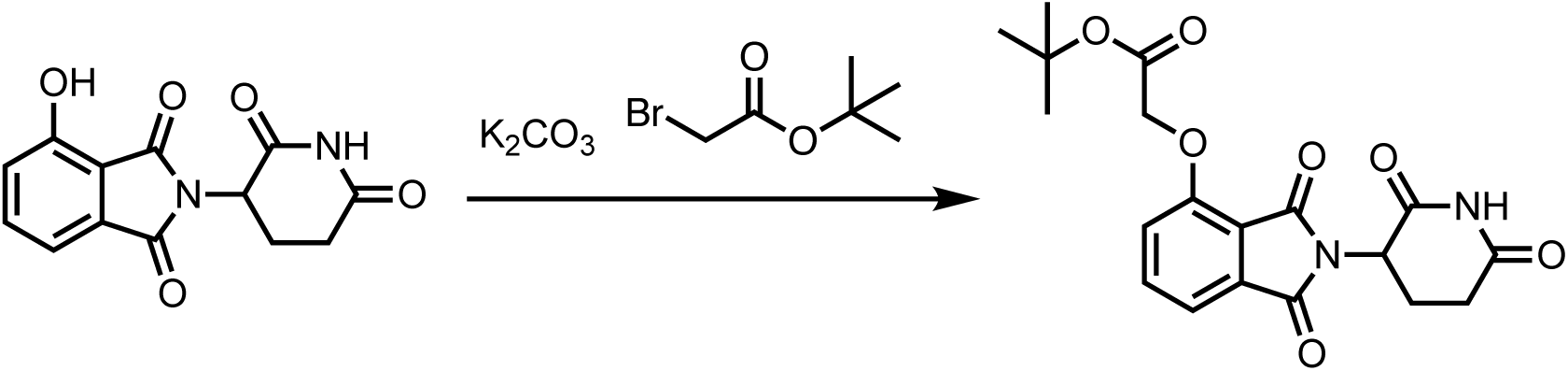

According to a reported literature^59^, 2-(2,6-dioxopiperidin-3-yl)-4-hydroxyisoindoline-1,3-dione (1 equiv) was dissolved in DMF at room temperature before the addition of K_2_CO_3_ (1.5 equiv) and tert-butyl bromoacetate (1.0 equiv). After 2 hours, the mixture was charged with DI H_2_O/EtOAc. The organic layer was dried over anhydrous MgSO_4_, concentrated under reduced pressure, and purified via flash column chromatography, which provided the title compound.

***tert*-Butyl (1-((2-(2,6-dioxopiperidin-3-yl)-1,3-dioxoisoindolin-4-yl)oxy)-2-oxo-6,9,12-trioxa-3-azatetradecan-14-yl)carbamate**

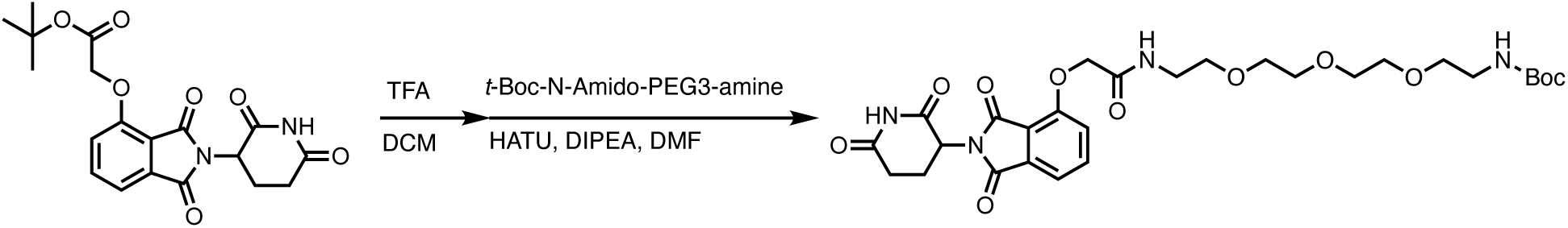

To the Boc-protected intermediate (1.0 equiv) in DCM, TFA (2.0 equiv) was added. The resulting solution was stirred for 1 hour to fully remove the Boc group, indicated by UPLC. The solvent was then removed under reduced pressure before re-dissolving in dry DMF. t-Boc-*N*-amido-PEG3-amine (1.1 equiv), HATU (1.5 equiv), and DIPEA (3.0 equiv) were successively charged to the reaction solution. After 2 hours, the reaction was charged with DI H_2_O/EtOAc. The organic layer was dried over anhydrous MgSO_4_, concentrated under reduced pressure, and purified via flash column chromatography, which provided the title compound.

***N*-(5-(3,5-Difluorobenzyl)-1*H*-indazol-3-yl)-4-(4-(17-((2-(2,6-dioxopiperidin-3-yl)-1,3-dioxoisoindolin-4-yl)oxy)-2,16-dioxo-6,9,12-trioxa-3,15-diazaheptadecyl)piperazin-1-yl)-2-((tetrahydro-2*H*-pyran-4-yl)amino)benzamide**

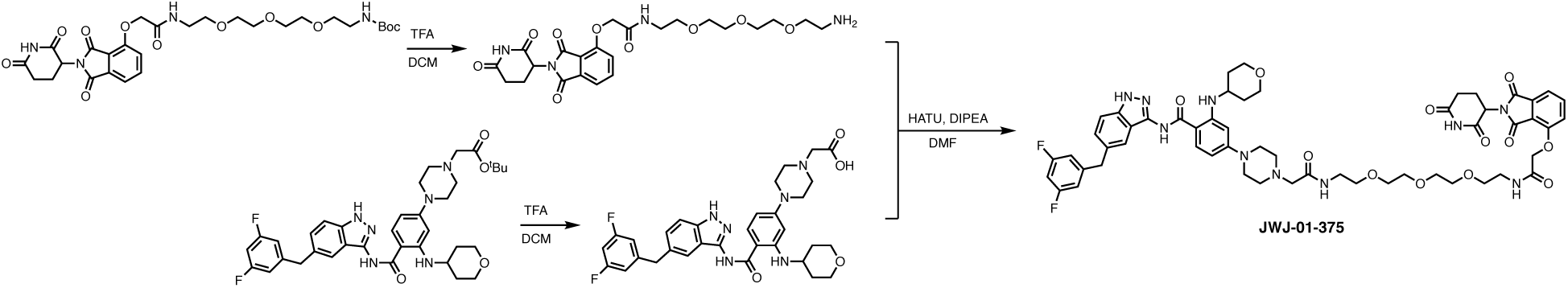

To the Boc-protected intermediate (1.0 equiv) in DCM, TFA (2.0 equiv) was added. The resulting solution was stirred for 1 hour to fully remove the Boc group, indicated by UPLC. The solvent was then removed under reduced pressure before re-dissolving in dry DMF. Meanwhile, the *tert*-butyl ester was dissolved in DCM before the addition of TFA (3.0 equiv). The solution was stirred for 5 hours to fully remove the ester group, monitored by UPLC. The solvent as well as excess TFA were evaporated under reduced pressure. The freshly generated acid (1.0 equiv) in DMF, HATU (1.5 equiv), and DIPEA (3.0 equiv) were successively added to Boc-deprotected amine solution. Upon the completion of the amide coupling reaction indicated by UPLC, the residue was purified by prep HPLC to yield the desired product (8.1 mg, 17%) with 94% purity. ^1^H NMR (500 MHz, DMSO-*d*_6_) δ 12.70 (s, 1H), 11.13 (s, 1H), 10.18 (s, 1H), 8.68 (t, *J* = 5.6 Hz, 1H), 8.01 (t, *J* = 5.8 Hz, 1H), 7.82 (dd, *J* = 17.1, 8.7 Hz, 2H), 7.51 (d, *J* = 7.3 Hz, 1H), 7.46 (s, 1H), 7.41 (dd, *J* = 8.6, 5.9 Hz, 2H), 7.26 (d, *J* = 8.6 Hz, 1H), 7.04–6.94 (m, 3H), 6.27 (d, *J* = 8.8 Hz, 1H), 6.18 (s, 1H), 5.12 (dd, *J* = 12.8, 5.4 Hz, 1H), 4.79 (s, 2H), 4.03 (d, *J* = 9.1 Hz, 8H), 3.81 (dd, *J* = 9.4, 5.6 Hz, 4H), 3.52 (s, 10H), 3.47 (d, *J* = 5.7 Hz, 4H), 3.32 (q, *J* = 5.7 Hz, 4H), 3.22 (s, 4H), 2.90 (ddd, *J* = 17.6, 13.6, 5.4 Hz, 1H), 2.65–2.51 (m, 2H), 2.08–2.01 (m, 1H), 1.93 (d, *J* = 12.7 Hz, 2H), 1.34 (q, *J* = 13.0, 11.2 Hz, 2H). MS (ESI) m/z 1093.6 (M + H)^+^.

***N*-(5-(3,5-Difluorobenzyl)-1*H*-indazol-3-yl)-4-(4-(14-((2-(2,6-dioxopiperidin-3-yl)-1,3-dioxoisoindolin-4-yl)oxy)-2,13-dioxo-6,9-dioxa-3,12-diazatetradecyl)piperazin-1-yl)-2-((tetrahydro-2*H*-pyran-4-yl)amino)benzamide**

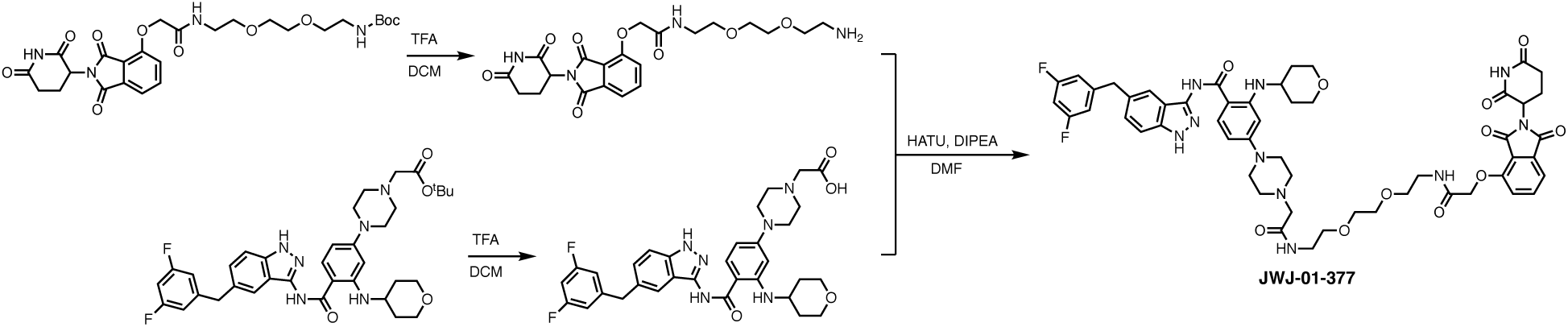

By employment of the similar procedure to JWJ-01-375, from the corresponding carboxylic acid (1.0 equiv), the title compound was prepared (6.7 mg, 17%) with 99% purity. ^1^H NMR (500 MHz, DMSO-*d*_6_) δ 12.70 (s, 1H), 11.13 (s, 1H), 10.17 (s, 1H), 8.68 (s, 1H), 8.02 (s, 1H), 7.87–7.79 (m, 2H), 7.51 (d, *J* = 7.3 Hz, 1H), 7.46 (s, 1H), 7.41 (d, *J* = 8.6 Hz, 2H), 7.28 (s, 1H), 7.05–6.96 (m, 3H), 6.29 (s, 1H), 6.18 (s, 1H), 5.12 (dd, *J* = 12.8, 5.4 Hz, 2H), 4.80 (s, 2H), 4.03 (d, *J* = 8.0 Hz, 6H), 3.82 (d, *J* = 11.2 Hz, 2H), 3.69 (s, 1H), 3.57–3.52 (m, 5H), 3.47 (td, *J* = 5.7, 2.3 Hz, 7H), 3.33 (q, *J* = 5.5 Hz, 4H), 3.22 (s, 4H), 2.94–2.86 (m, 1H), 2.63–2.53 (m, 2H), 2.07–2.00 (m, 1H), 1.93 (d, *J* = 12.3 Hz, 2H), 1.34 (d, *J* = 11.0 Hz, 2H). MS (ESI) m/z 1049.7 (M + H)^+^.

***N*-(5-(3,5-Difluorobenzyl)-1*H*-indazol-3-yl)-4-(4-(2-((8-(2-((2-(2,6-dioxopiperidin-3-yl)-1,3-dioxoisoindolin-4-yl)oxy)acetamido)octyl)amino)-2-oxoethyl)piperazin-1-yl)-2-((tetrahydro-2*H*-pyran-4-yl)amino)benzamide**

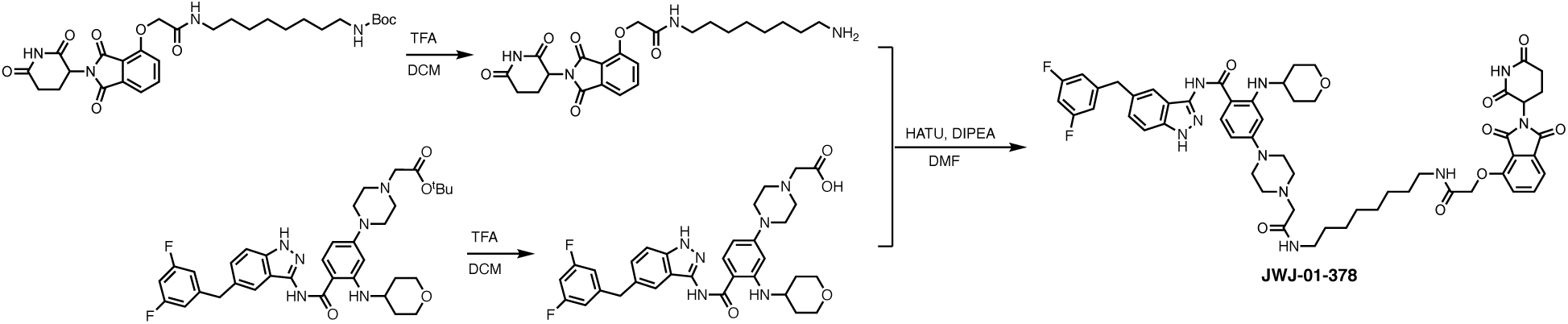

By employment of the similar procedure to JWJ-01-375, from the corresponding carboxylic acid (1.0 equiv), the title compound was prepared (6.3 mg, 16%) with 92% purity. ^1^H NMR (500 MHz, DMSO-*d*_6_) δ 12.69 (s, 1H), 11.13 (s, 1H), 10.18 (s, 1H), 8.54 (t, *J* = 5.7 Hz, 1H), 7.95 (t, *J* = 5.8 Hz, 1H), 7.87–7.77 (m, 2H), 7.51 (d, *J* = 7.3 Hz, 1H), 7.46 (s, 1H), 7.40 (t, *J* = 7.8 Hz, 2H), 7.27 (s, 1H), 7.05–6.95 (m, 3H), 6.28 (d, *J* = 8.7 Hz, 1H), 6.19 (s, 1H), 5.12 (dd, *J* = 12.8, 5.4 Hz, 1H), 4.77 (s, 2H), 4.02 (d, *J* = 18.5 Hz, 6H), 3.90–3.80 (m, 4H), 3.48 (t, *J* = 10.8 Hz, 4H), 3.32–3.06 (m, 8H), 2.96–2.88 (m, 1H), 2.64–2.51 (m, 2H), 2.07–2.01 (m, 1H), 1.93 (d, *J* = 12.4 Hz, 2H), 1.44 (t, *J* = 6.9 Hz, 4H), 1.34 (d, *J* = 10.6 Hz, 2H), 1.26 (s, 8H). MS (ESI) m/z 1045.7 (M + H)^+^.

***N*-(5-(3,5-Difluorobenzyl)-1*H*-indazol-3-yl)-4-(4-(2-((8-(2-((2-(1-methyl-2,6-dioxopiperidin-3-yl)-1,3-dioxoisoindolin-4-yl)oxy)acetamido)octyl)amino)-2-oxoethyl)piperazin-1-yl)-2-((tetrahydro-2*H*-pyran-4-yl)amino)benzamide**

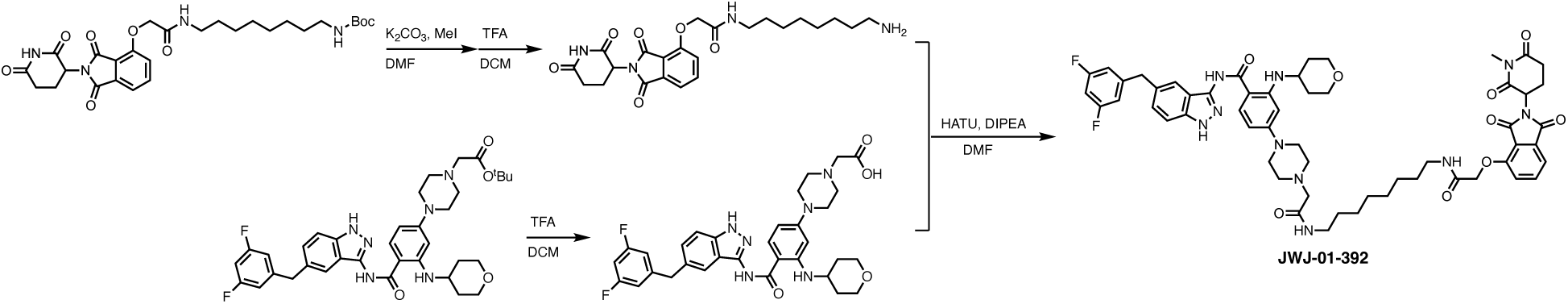

By employment of the similar procedure to JWJ-01-375, from the corresponding carboxylic acid (1.0 equiv), the title compound was prepared (6.9 mg, 18%) with 99% purity. ^1^H NMR (500 MHz, DMSO-*d*_6_) δ 12.71 (s, 1H), 10.19 (s, 1H), 8.55 (d, *J* = 5.8 Hz, 1H), 7.97 (t, *J* = 5.9 Hz, 1H), 7.88–7.78 (m, 2H), 7.51 (d, *J* = 7.2 Hz, 1H), 7.46 (s, 1H), 7.40 (dd, *J* = 8.6, 6.0 Hz, 2H), 7.26 (d, *J* = 8.5 Hz, 1H), 7.00 (dd, *J* = 20.8, 8.7 Hz, 3H), 6.27 (d, *J* = 9.0 Hz, 1H), 6.18 (s, 1H), 5.19 (dd, *J* = 13.0, 5.4 Hz, 1H), 4.77 (s, 2H), 4.02 (d, *J* = 15.0 Hz, 6H), 3.82 (d, *J* = 8.4 Hz, 4H), 3.69 (s, 2H), 3.48 (t, *J* = 10.8 Hz, 4H), 3.21 (s, 2H), 3.14 (q, *J* = 6.6 Hz, 4H), 3.02 (s, 3H), 2.97–2.91 (m, 1H), 2.80– 2.74 (m, 1H), 2.55 (d, *J* = 8.5 Hz, 1H), 2.05 (d, *J* = 12.7 Hz, 1H), 1.93 (d, *J* = 12.5 Hz, 2H), 1.43 (t, *J* = 6.9 Hz, 4H), 1.34 (d, *J* = 10.4 Hz, 2H), 1.26 (s, 8H). MS (ESI) m/z 1059.9 (M + H)^+^.

### Additional compounds

Entrectinib (Selleckchem, #S7998), GNF-8625 (MedChemExpress, #HY-131706A), CG-428 (Tocris, #7425), carfilzomib (MedChemExpress, #HY-10455), MLN4924 (Calbiochem, #5054770001), lenalidomide (Millipore Sigma, #901558) and bafilomycin A1 (Cambridge Bioscience, # b0025) were purchased from the indicated sources.

### Cell line culturing

KM12 (kindly provided by the MD Anderson Cancer Center, Houston, TX) and HEL 92.1.7 (ATCC, #TIB-180) cells were cultured in RPMI 1640 medium supplemented with 10% fetal bovine serum, 100 U/mL penicillin, and 100 μg/mL streptomycin at 37°C with 5% CO_2_. DF-1 cells (ATCC, #CRL-12203) were cultured in Dulbecco’s modified Eagle medium (DMEM) supplemented with 10% fetal bovine serum, 100 U/mL penicillin, and 100 μg/mL streptomycin at 39 °C with 5% CO_2_. NIH/3T3-tv-a cells^60^ were cultured in DMEM supplemented with 10% calf serum, 100 U/mL penicillin, and 100 μg/mL streptomycin at 37°C with 5% CO_2_. Cell lines were confirmed for mycoplasma negative monthly using the MycoProbe Mycoplasma detection kit (R&D systems, #CUL001B).

### Viral production and stable cell line generation

For the generation of NIH/3T3 cells expressing TRK fusions, DF-1 cells were transfected with the indicated RCAS plasmid using X-tremeGENE 9 DNA transfection reagent (Roche, #XTG9-RO) according to the manufacturer’s protocol. Viral supernatant from DF-1 cultures was sterile-filtered and added to NIH/3T3-tv-a cells every eight hours for three cycles.

### Immunoblotting

Immunoblotting was performed as previously described^61^. Cell lines were lysed in RIPA buffer containing cOmplete protease inhibitor (Millipore Sigma, #11873580001), PhosSTOP phosphatase inhibitor (Millipore Sigma, #4906837001), and 0.1% Benzonase (Millipore Sigma, # 712063) on the ice for 1 hour. Lysates were centrifuged at 12000 rpm for 10 minutes at 4°C and clarified supernatants were transferred to the pre-chilled tubes. Protein lysates were quantified and loaded in equal amounts on 4-12% SDS-PAGE gradient gels, followed by transfer onto nitrocellulose membranes using an iBlot3 (Thermo Fisher). The membranes were incubated overnight with the indicated primary antibodies: pan-TRK (Cell Signaling Technology, #9299S), β-Actin (Cell Signaling Technology, #8H10D10), p44/42 MAPK (ERK1/2) (Cell Signaling Technology, #4696), Phospho-p44/42 MAPK (ERK1/2) (Thr202/Tyr204) (Cell Signaling Technology, #4370), and ALK (Cell Signaling Technology, #3633). Membranes were washed the next day with TBS-T (TBS with 0.1% Tween), incubated with fluorescently labeled infrared secondary antibodies (Li-COR) at room temperature for 1 hour, and imaged using the LI-COR Odyssey® Imaging System.

### Analysis of cell viability in 2D-adherent monolayer and 3D-spheroid culture

2D-adherent monolayer and 3D-spheroid culture experiments were performed as previously described^32, 61^ with minor modifications. For 2D-adherent viability assays, KM12 cells were plated in white 384-well culture plates (Corning, #3570) at a density of 500 cells/well in 50 μL media. HEL cells, which are suspension cultures, were also plated in white 384-well culture plates (Corning, #3570) at a density of 1000 cells/well in 50 µL media. For ultra-low adherent 3D-spheroid viability assays, KM12 cells were plated in PrimeSurface 384-well 3D culture spheroid plates (S-bio, #MS-9384WZ) at a density of 500 cells/well in 50 µL media. The next day, cells were treated with compounds using a D300e digital dispenser (HP), and all wells were normalized to 100 nL total DMSO. For the drug efflux inhibitor experiments, cells were treated simultaneously with test compounds and zosuquidar 3HCl (Selleck Chem, #S1481) or tariquidar (MedChemExpress, #HY-10550) and incubated for 120 hours. For experiments where a second dose of compound was administered, cells were treated with indicated compounds again at 72 hours. In all experiments, after 120 hours from the start of treatment, cells were equilibrated at room temperature for 30 minutes, followed by the addition of 10 µL CellTiter-Glo (Promega, #G7570) per well, for 15 minutes with shaking at room temperature. Luminescence was measured on the CLARIOstar Plus Plate Reader (BMG LabTech). The data was normalized to DMSO-treated control wells and analyzed using GraphPad PRISM v10.

### Data-independent acquisition (DIA)-based quantitative proteomics

#### Cell culture and compound treatment

KM12 cells were plated at a density of 40000 cells/well in 200 µL of media in 96 well plates (Greiner, #655180) using a Viaflo liquid handler (Integra). After 24 hours, cells were treated with compounds using drug dispenser D300e (Tecan) in triplicate. Cells were treated using Integra Viaflow by transferring media off of cells, dispensing and mixing on a compound plate to resuspend them, and then transferring the media plus drug back onto cells prior to incubation. Cells were treated with 100 nM of JWJ-01-378 for two hours followed by protein fractionation, digestion, and mass spectrometry DIA analysis.

#### Protein fractionation or whole cell lysis

Following compound incubation, the treated cells were dissociated with Accutase enzyme (Sigma-Aldrich, #A6964) and washed with PBS. For whole cell lysis, cells were resuspended with a buffer composed of 10 mM Tris with 15 mM NaCl, 60 mM KCl (pH 8.0), and benzonase, followed by incubation at 37 °C for 15 minutes prior to protein digestion.

#### Protein digestion and peptide preparation for mass spectrometry acquisition

Proteins were denatured by heating to 95 °C for 5 minutes. Dithiothreitol (DTT) was added to plates using a Combi reagent dispenser (Thermo Fisher) to a final concentration of 4 mM DTT in-solution. Proteins were incubated in DTT solution at 60 °C for 30 minutes to reduce proteins. Iodoacetamide (IAM) was added using a Combi reagent dispenser to a final concentration of 20 mM in-solution. Proteins were incubated with IAM for 30 minutes in the dark at room temperature to perform alkylation. The Combi reagent dispenser was used to dispense DTT to all samples again to quench alkylation reaction. Pre-washed carboxyl beads were bound to proteins, and then 100% ethanol was added via a Combi reagent dispenser to precipitate proteins and facilitate binding onto magnetic carboxyl-coated beads (Cytiva). Proteins bound to carboxyl beads were washed five times in 50% ethanol via KingFisher Apex and then eluted into Promega Sequencing Grade Trypsin at a 1:10 enzyme to protein ratio. Proteins were digested into peptides for approximately 16-20 hours with trypsin at 37 °C.

#### Mass spectrometry preparation and acquisition

After digestion, trypsin was quenched with 10% formic acid using an Opentron 2 (OT2) liquid handling robot. Samples were transferred off of beads and diluted 6-fold prior to EvoTip loading, using OT2 liquid handling robot^62^. Approximately 100 ng of peptide was loaded onto EvoTips for data-independent acquisition. Both sample sets were randomized across their condition groups and acquired on timsTOF Ultra 2 coupled to EvoSep liquid chromatography system over a 22 minute gradient, equivalent to 60 samples per day.

#### Mass spectrometry data analysis

Raw mass spectrometry data were processed using DIANN v1.8.1 within the quantms NextFlow pipeline^63–65^, which performed peptide and protein identification and quantification of DIA LC-MS data. Data quality control involved assessing protein coefficient of variation (CV), quantifying missing values per sample replicate and protein, and evaluating protein detection depth to ensure sufficient protein identification coverage. For each condition, log_2_-transformation, scaled, and normalized data was analyzed using a linear modeling framework implemented in the limma package. The resulting data were subjected to a moderated *t*-test to assess statistical significance also implemented in the limma package. For protein quantification, the two most abundant peptides per protein under DMSO conditions were selected, applying a minimum threshold of 2^14 raw peptide intensity value. Data was plotted and analyzed using ggplot2 in R. The raw proteomics datasets generated during this study are available at the ProteomeXchange Consortium via the PRIDE partner repository^56, 57^ with the dataset identifiers PXD065035.

