## Supplementary Information for "Development of PROTACs for targeted degradation of oncogenic TRK fusions"

[‡] Equal contribution

***Correspondence should be addressed to:**

**Supplementary Tables**

| Compound | 2D-adherent monolayer $IC_{50}$ (nM)[a] | 3D-spheroid suspension $IC_{50}$ (nM)[a] |
|---|---|---|
| Entrectinib | 0.7 | 2.0 |
| JWJ-01-375 | 22.7 | 31.6 |
| JWJ-01-377 | 10.8 | 26.4 |
| JWJ-01-378 | 6.2 | 4.2 |
| JWJ-01-392 | 86.7 | 109.7 |
| GNF-8625 | 2.5 | 5.9 |
| CG-428 | 0.9 | 1.0 |
| [a]$IC_{50}$ values are associated with data in Figure 4. | | |

**Table S1 |** Antiproliferative effects of TRK inhibitors and PROTACs in KM12 cells cultured as 2D-adherent monolayers or 3D-spheroid suspensions.

### Supplementary Figures

A

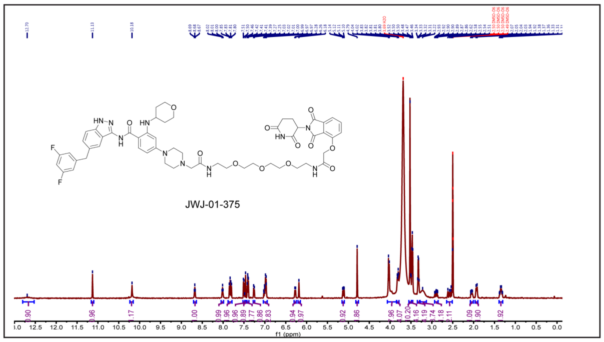


B

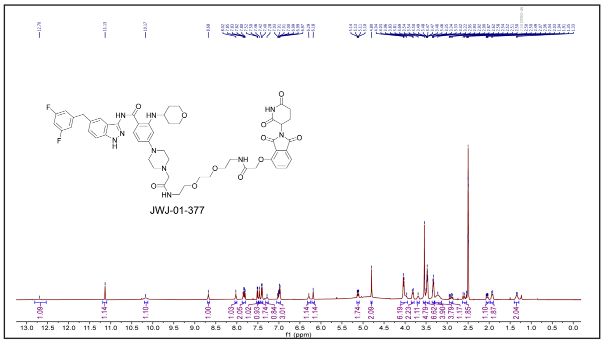


C

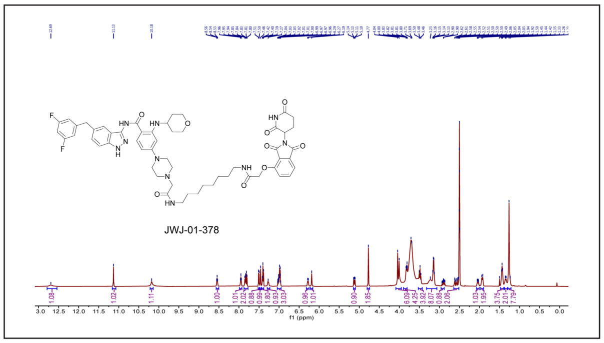


D

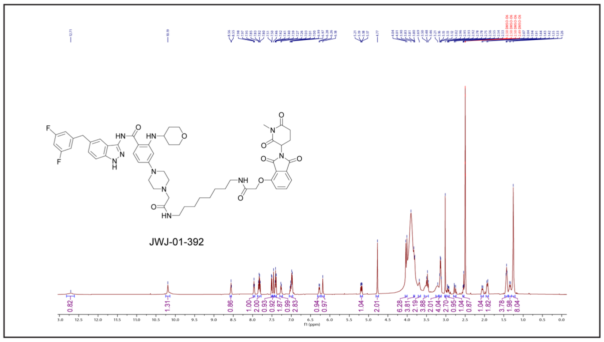


**Figure S1 | NMR spectra for the indicated compounds.** (A) $^{1}$H NMR (500 MHz, DMSO-*d6*) spectra of JWJ-01-375. (B) $^{1}$H NMR (500 MHz, DMSO-*d6*) spectra of JWJ-01-377. (C) $^{1}$H NMR (500 MHz, DMSO-*d6*) spectra of JWJ-01-378. (D) $^{1}$H NMR (500 MHz, DMSO-*d6*) spectra of JWJ-01-392.

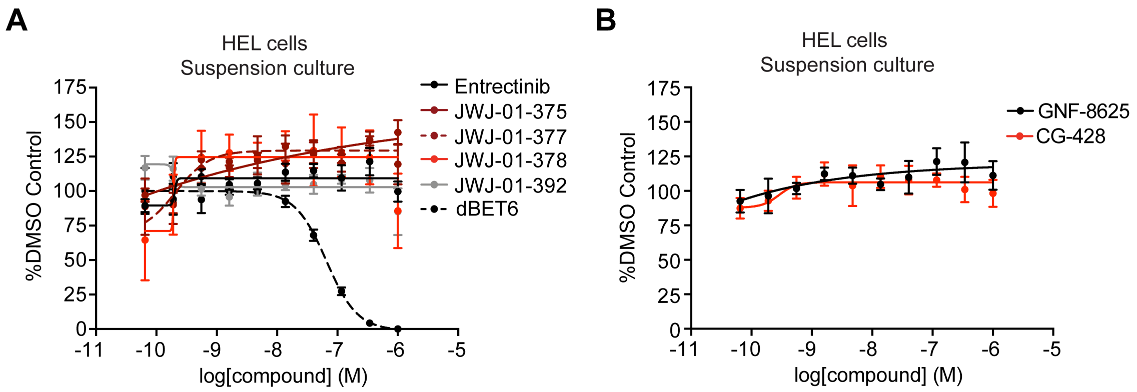


**Figure S2 | TRK-targeting compounds do not impact HEL cell viability.** (A-B) DMSO-normalized antiproliferation of HEL cells cultured in suspension and treated with indicated compounds for 120 hours. Data are presented as mean ± s.d. of *n* = 4 biologically independent samples and are representative of *n* = 3 independent experiments.

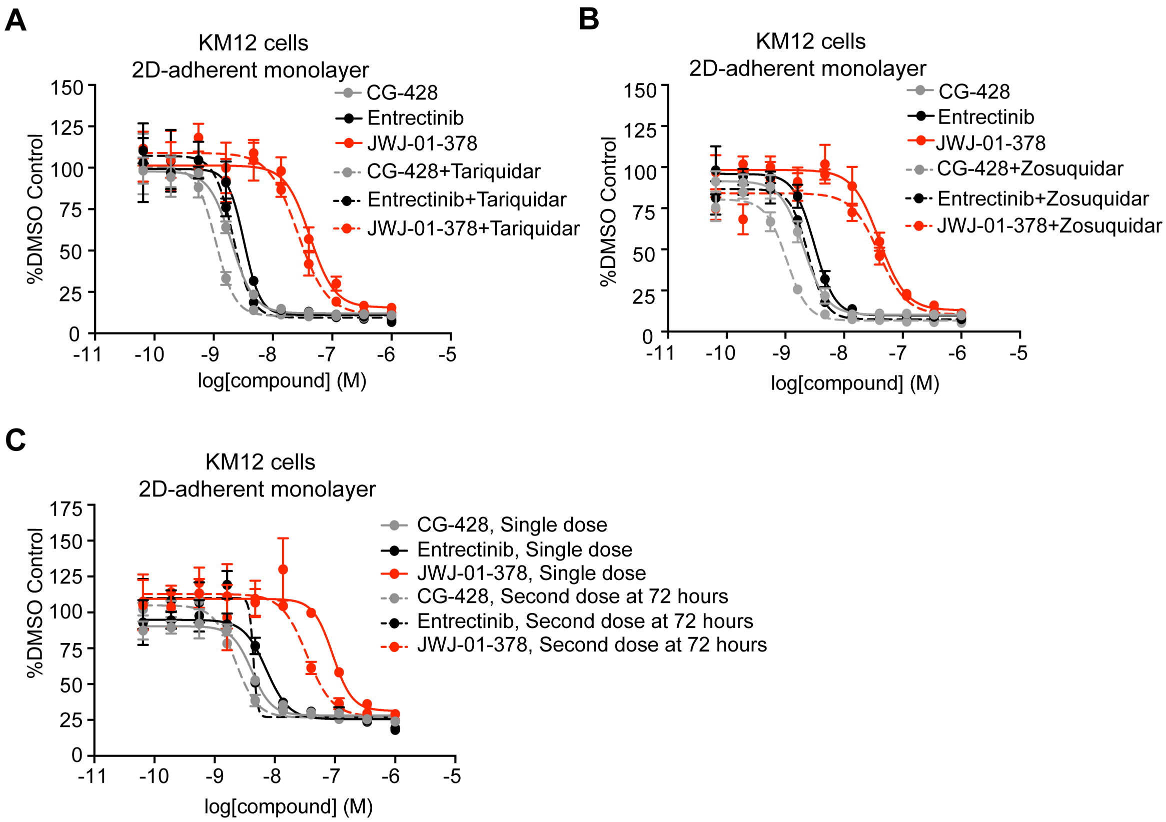


**Fig. S3 | Drug efflux inhibitors or an additional administration of compound do not improve the biological responses of TRK-targeting inhibitors or PROTACs.** (A) DMSO-normalized antiproliferation of KM12 cells cultured as 2D-adherent monolayers and treated with indicated compounds for 120 hours. (B) DMSO-normalized antiproliferation of KM12 cells cultured as 2D-adherent monolayers and treated with indicated compounds for 120 hours. (C) DMSO-normalized antiproliferation of KM12 cells cultured as 2D-adherent monolayers and treated with indicated compounds for a single dose for 120 hours or a second dose at 72 hours before evaluation at 120 hours. Data in (A-C) are presented as mean ± s.d. of $n$ = 4 biologically independent samples and are representative of $n$ = 3 independent experiments.
